# Illuminating the Role of Asymmetric Mitochondrial Fission on Beta-Cell Health

**DOI:** 10.1101/2025.11.25.690461

**Authors:** Philipp Henning, Julia Schultz, Simone Baltrusch, Adelinde M. Uhrmacher

## Abstract

Mitochondrial dynamics play a critical role in the development of aging-related diseases such as type 2 diabetes mellitus.

To investigate how mitochondrial dynamics influence cellular behavior in pancreatic beta-cells, we developed a rule-based, multi-level simulation model of insulin secretion. The pancreatic beta cell model encompasses metabolic pathways (glycolysis and oxidative phosphorylation), compartmental processes (mitochondrial fusion and fission), and cellular processes (insulin secretion), allowing for the investigation of their interplay. The rule-based simulation model captures the high plasticity of these organelles and integrates and builds upon insights from various experimental studies and previous simulation models. Its rule-based specification facilitates the exploration of new hypotheses, the integration of new knowledge and data, and the successive extension of the model.

The results of our simulation experiments underscore the importance of peripheral, sorted mitochondrial fission in maintaining mitochondrial health. Downregulation of the fission-associated anchor proteins Fis1 and MFF impacts mitochondrial structure and function differently, highlighting their distinct roles in maintaining mitochondrial health and cellular biogenesis, respectively. With respect to insulin secretion, Drp1 suppression shows that beta cells become unresponsive to glucose, whereas Fis1 downregulation only attenuates the cellular response. The simulation model and simulation results corroborates experimental findings and contribute to a deeper understanding of the mechanisms involved in mitochondrial dynamics of pancreatic beta cells and their relation to metabolic dysregulation in type 2 diabetes mellitus.

**Author summary:** Mitochondria, often described as the powerhouses of the cell, undergo constant changes through fusion and fission processes. These dynamics are essential for maintaining cell health and proper functioning. In type 2 diabetes mellitus, this balance can be disrupted. In this work, we developed a multi-level, rule-based simulation model to analyze processes of mitochondrial dynamics and their impact on insulin secretion within pancreatic beta cells. Our model captures diverse biological processes that operate at different but interconnected organizational levels, including energy metabolism, mitochondrial dynamics, and insulin secretion. We found that peripheral fission plays a crucial role in determining whether cells secrete insulin properly. In addition, downregulation of the fission-associated proteins MFF and Fis1 reveals a distinct impact on the structure and function of the mitochondrial network, as well as on insulin secretion in pancreatic beta cells. The simulation model and results provide insights into how mitochondrial dynamics affect beta cell metabolism and insulin release. It enables the study of dynamics at different organizational levels, and its rule-based approach facilitates the integration of new knowledge (e.g., by updating or adding specific rules) and experimental data.

## 1 Introduction

Mitochondria play a central role in health regulation and disease progression [1]. The dysfunction of mitochondria is associated with various diseases, including neurodegenerative disorders, cardiovascular diseases, cancer, and diabetes. The development of type 2 diabetes mellitus is often associated with pancreatic beta-cell exhaustion due to peripheral insulin resistance. Under physiological conditions, pancreatic beta-cells sense glucose to adjust insulin exocytosis and maintain glucose homeostasis. In this process, mitochondria play a vital role. An increased glucose level leads to the production of metabolites, in particular, ATP (partly within the mitochondria and partly via cytosolic glycolysis) that operate jointly with cytosolic Ca^2+^ to stimulate insulin secretion. Fission and fusion processes shape the mitochondrial structure, allowing for the selective removal of damaged or dysfunctional mitochondria and, consequently, effective mitochondrial quality control. Mitochondrial dysfunction, on the other hand, appears to be critical in the development of type 2 diabetes [2]. Figure 1 illustrates the fragmented mitochondrial structure within beta cells of type 2 diabetes mellitus patients.

**Fig 1.**
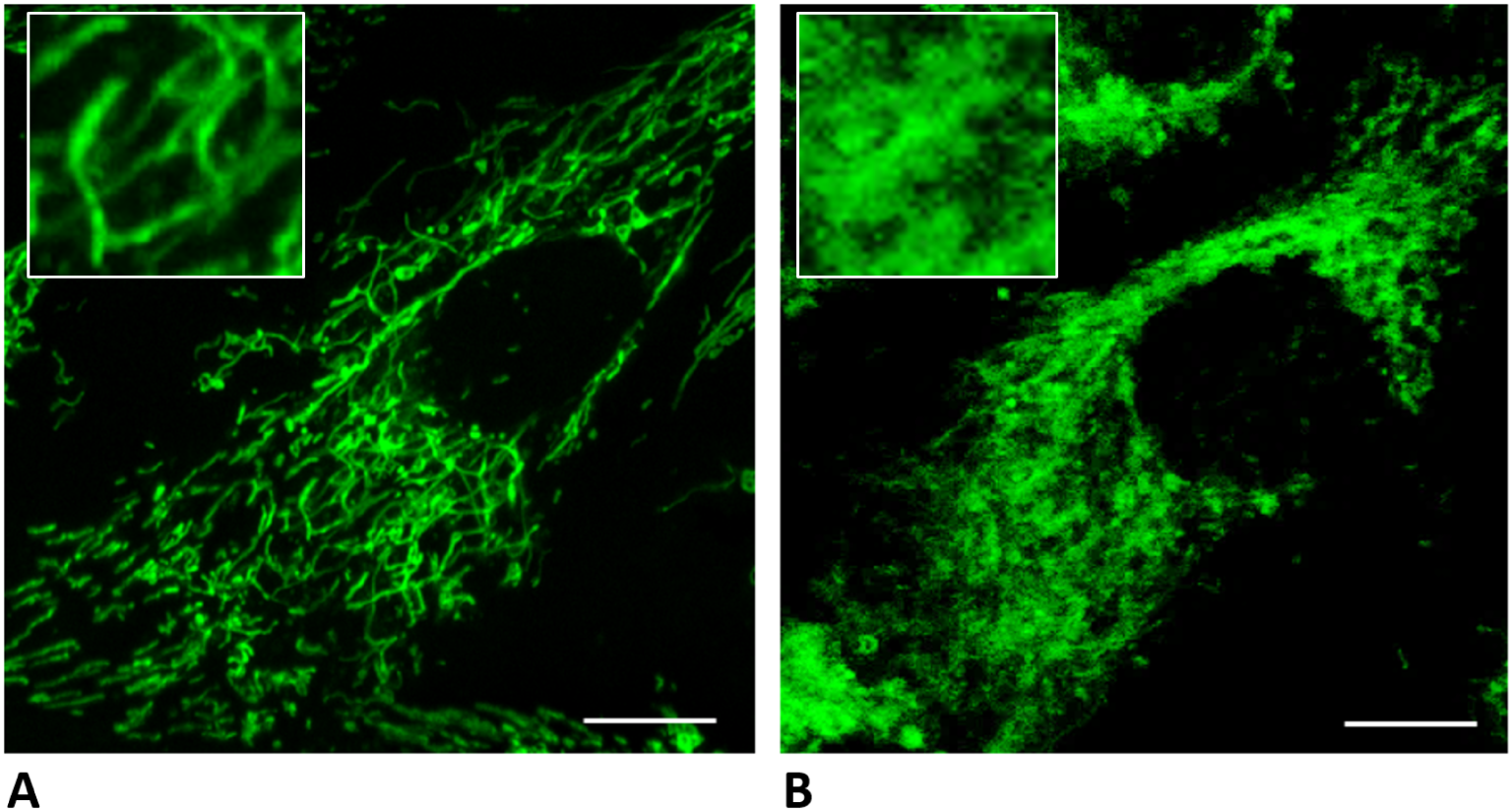
Mitochondrial network visualized with MitoTrackerGreen in an isolated human pancreatic beta cell from a donor without (A) and with (B) type 2 diabetes mellitus. Scale bar: 10 µm

Our goal is to develop a simulation model that captures the interplay between intracellular and mitochondrial metabolism, mitochondrial dynamics, and the insulin secretion pathway, thereby furthering our understanding of the response of pancreatic beta cells to glucose and the subsequent impact of mitochondrial dysregulation on disease manifestation and progression.

In order to understand these mechanics, various simulation models have already been developed. However, their focus is either a) on the secretion of insulin, b) intracellular and mitochondrial metabolism, or c) the mitochondrial network dynamics.

a) Ordinary differential equation (ODE)-based models of calcium signaling and glucose-stimulated insulin secretion (GSIS) in beta cells have been reviewed by Félix-Martínez and Godínez-Ferńandez [3]. The degrees of complexity vary from simulation models that only include calcium signaling to those that include rudimentary ATP dynamics and several ion fluxes. However, all these simulation models investigate the triggering pathway and, thus, only capture the initial seconds to minutes of the calcium and GSIS dynamics, focusing on the oscillation of the mitochondrial membrane potential and the concentration of Ca^2+^. Other models [4, 5] not only capture the triggering but also the amplifying pathway of insulin secretion and can simulate the insulin secretion for several hours.
b) The metabolic models of interest for the insulin secretion pathway are those that focus on glucose metabolism, specifically glycolysis and oxidative phosphorylation (oxPhos). The glycolysis is modeled in great detail in [6, 7], where each pathway enzyme is characterized and simulated. ODE models of oxidative phosphorylation can be found in [8–10], where, similar to the glycolysis models, the key enzymes of the pathway are simulated. The models are either used to study the steady-state of the system or the dynamics over several minutes to an hour.
c) A wide range of models with different modeling approaches is employed regarding mitochondrial fission-fusion dynamics. This includes stochastic models with a spatial component [11–14], ODE models [15], and graph-based models [16]. These models are used to investigate different aspects of the mitochondrial network dynamics, like its self-organization [16], damage regulation [12, 13, 15], coupling to microtubules [14], or propagation of mutations [11]. As changes in the mitochondrial network are slow in comparison to metabolic processes, these models cover the dynamic over several hours to days.

None of the above simulation models captures dynamics at multiple levels of organization, i.e., intra-mitochondrial, mitochondrial, intracellular, and cellular, that we deem important to analyze the interdependence between mitochondrial dynamics and the response of beta-cells to glucose. In the following, we exploit a multi-level rule-based modeling and simulation approach [17] to develop a beta-cell model that integrates the secretion of insulin, intracellular and mitochondrial metabolism, and mitochondrial network dynamics.

## 2 Materials and methods

### 2.1 Simulation Tool

We used ML-Rules3 [18] as our modeling and simulation environment. It constitutes a Rust implementation of the multi-level rule-based modeling and simulation approach introduced in [17]. Its semantics is basically a Continuous Time Markov Chain, being formally specified in [19]. ML-Rules3 is a rule-based stochastic simulator that supports dynamic compartmentalization, allowing particles to be nested within compartments and moved between them during simulations, and the merging and division of compartments.

The rule-based modeling approach enables a compact notation inspired by chemical reaction notation. For example, the reaction of particles A and B to particle C with the rate *k*_1_ is written as:

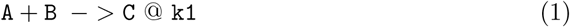

Here, a single @ denotes that the reaction rate follows the mass-action law with coefficient k1. Using a double symbol (@@) instead applies the specified rate directly, bypassing mass-action assumptions, allowing the use of a wide range of rate equations. For example, the fermentation of pyruvate (simulated by its removal) shown below follows Michaelis-Menten kinetics.

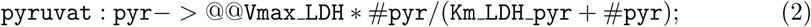

The rate specifies an exponentially distributed sojourn time between the events. Further properties of a particle, like the volume or phosphorylation state, can be added by using attributes, which are noted after the species in brackets. These attributes can be used to filter participants in reactions, modify rate expressions, and can be updated during simulation. The use of a stochastic approach is essential due to the low concentrations of certain particles. For instance, a Min6 cell (a mouse insulinoma cell line with properties similar to a primary beta cell) typically contains fewer than 100 mitochondria [20], each displaying only a few spots per mitochondrion where fission proteins may attach [21, 22]. In such cases, stochastic fluctuations can significantly influence system behavior. At the same time, the stochastic approach is not ideal for particles that are present in large numbers and undergo rapid reactions, like ATP. A single cell can contain more than a billion ATP molecules [23] and consume more than 100 million ATP molecules per second [24]. To simulate such molecules efficiently with the current simulator (for the Rust implementation, so far no hybrid simulator exists [25]), we aggregate them into packets representing many molecules, e.g., ATP 1000000 corresponds to one million ATP particles.

This abstraction maintains computational feasibility while preserving relevant dynamics. A key feature of ML-Rules3 is its support for nested particles and dynamic compartments, which enables the representation of cellular hierarchies and their changes during the model’s runtime. An example of nesting, denoted by curly brackets, is given in the initialization:

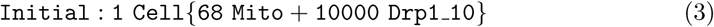

which describes that one cell contains 68 mitochondria and 10000 Drp1 clusters. Other simulation tools (e.g., Copasi [26] or VCell [27]) also allow the modeling of biological compartments, but do not offer the possibility to model fission, fusion, the generation or removal of compartments during runtime. A feature that is crucial to simulating the mitochondrial fission-fusion dynamics. The fission of a mitochondrion into two sister mitochondria can be written down by

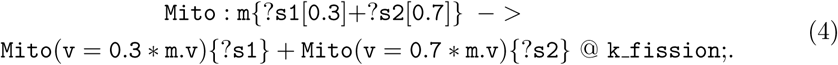

Here, the content nested in the Mito compartment, denoted by ?*s*, is divided unevenly (30% to 70%) between the two new ones, as well as the volume attribute v.

### 2.2 Parameter Estimation

Not all parameters required for the model can be directly obtained from the literature (see the Appendix 4.3 for a complete list). Some are estimated using inverse methods; these rely on the availability of data with which simulation results are compared [28]. In this study, we employed pyABC [29] and pySwarms [30], two Python-based tools for parameter inference. These tools facilitate the identification of model parameters that best replicate reference data. pyABC utilizes Approximate Bayesian Computation (ABC) to estimate the posterior distribution of a parameter. From this distribution, the parameter that most accurately reproduces the reference data can be selected. Additionally, the shape of the posterior distribution offers insights into parameter convergence, output sensitivity, and the potential existence of multiple plausible parameter values. However, ABC-based inference typically requires a substantial number of simulation runs (on the order of tens of thousands) to accurately estimate the distribution, making it feasible only for models with relatively short execution times. In contrast, pySwarms apply Particle Swarm Optimization (PSO), providing only the best-fitting parameter values without generating a posterior distribution. Since it requires fewer simulation runs, pySwarms is also suitable for models with longer execution times.

### 2.3 Sensitivity Analysis

To assess the influence of individual model parameters on a specific model output, we use the SALib Python library [31, 32] to conduct a variance-based global sensitivity analysis. Specifically, we applied Sobol’s method, which decomposes the output variance to quantify the contribution of each parameter and its interactions. The analysis yields first-order, second-order, and total-order sensitivity indices. The First-order indices measure the direct effect of a single parameter on the output variance. Second-order indices capture interactions between pairs of parameters, and total-order indices represent each parameter’s overall contribution, including its individual and interaction effects.

### 2.4 Cell Imaging

Human islets were donated by patients after pancreatic surgery and were used only with their prior consent. Islets were isolated by collagenase P (Roche Diagnostics, Mannheim, Germany) digestion and Ficoll gradient (Ficoll PM 400; Sigma, Seelze, Germany). Islets were seeded on glass-bottom MatTek dishes (MatTek Corporation, Ashland, MA, USA) and stained with 20 nmol/l MitoTracker^®^ Green FM (ThermoFisher Scientific, Waltham, MA, USA) for 30 minutes at 37°C. Mitochondrial morphology was analysed using a Fluoview FV10i confocal microscope (Olympus, Hamburg, Germany).

## 3 Simulation Model

Building a beta-cell model that captures both insulin secretion and the dynamics of mitochondrial fission and fusion is a difficult task due to the complexity of the involved processes. To simulate insulin secretion, the model must incorporate ATP generation and calcium flux, and to model the fission of mitochondria, the polymerization of the fission protein Drp1 needs to be included (see Sec. 3.1). We address this complexity by employing the divide-and-conquer approach and first construct three submodels: the fission-fusion model, the ATP-Insulin model, and the cAMP model, which we later integrate into the beta-cell model. These submodels are smaller and more manageable, allowing for the integration of behavior observed in the wet-lab and a less computationally expensive calibration to wet-lab data.

### 3.1 Submodels

#### Fission-Fusion Model

This submodel (see Fig. 2 top left) includes the binding and unbinding of Drp1 to mitochondrial anchor sites (which are later resolved into Fis1 and MFF), the fission and fusion of the mitochondria, and the (de)phosphorylation of Drp1. Due to technical limitations, Drp1 is simulated in clusters of 10 molecules (see Sec. 2.1). The central component of the model is the mitochondria, which are characterized by a volume attribute. Within the mitochondrial compartments, discrete anchor sites for Drp1 recruitment are nested. Unphosphorylated Drp1 can reversibly bind to these anchor sites [33]. The cAMP-dependent protein kinase (PKA) phosphorylates Drp1 if it is activated by cAMP [34], whereas Calcineurin (CaN) dephosphorylates Drp1 if it is activated by Ca^2+^ [35]. In this submodel, the concentrations of second messengers cAMP and Ca^2+^ are held constant. Fission events can occur when a sufficient number of Drp1 molecules are bound at an anchor site [36]. The fission rate itself depends on the size of the mitochondria, with larger mitochondria being more likely to split [37]. The fission event can either be a ”midzone” event, where two roughly equally large mitochondria are generated, or a ”peripheral” fission event, where one of the sister mitochondria is smaller. The data from [38] are used to draw normally distributed random numbers to realize the midzone and peripheral fission events. The fusion rule is independent of the size of the mitochondria [37] and uses a constant rate coefficient (see Appendix 4.3). To calibrate the fission and fusion dynamics, data for the Drp1 binding [36], frequency of fission events [37, 39], and number of mitochondria in a Min6 cell [20] were used.

**Fig 2.**
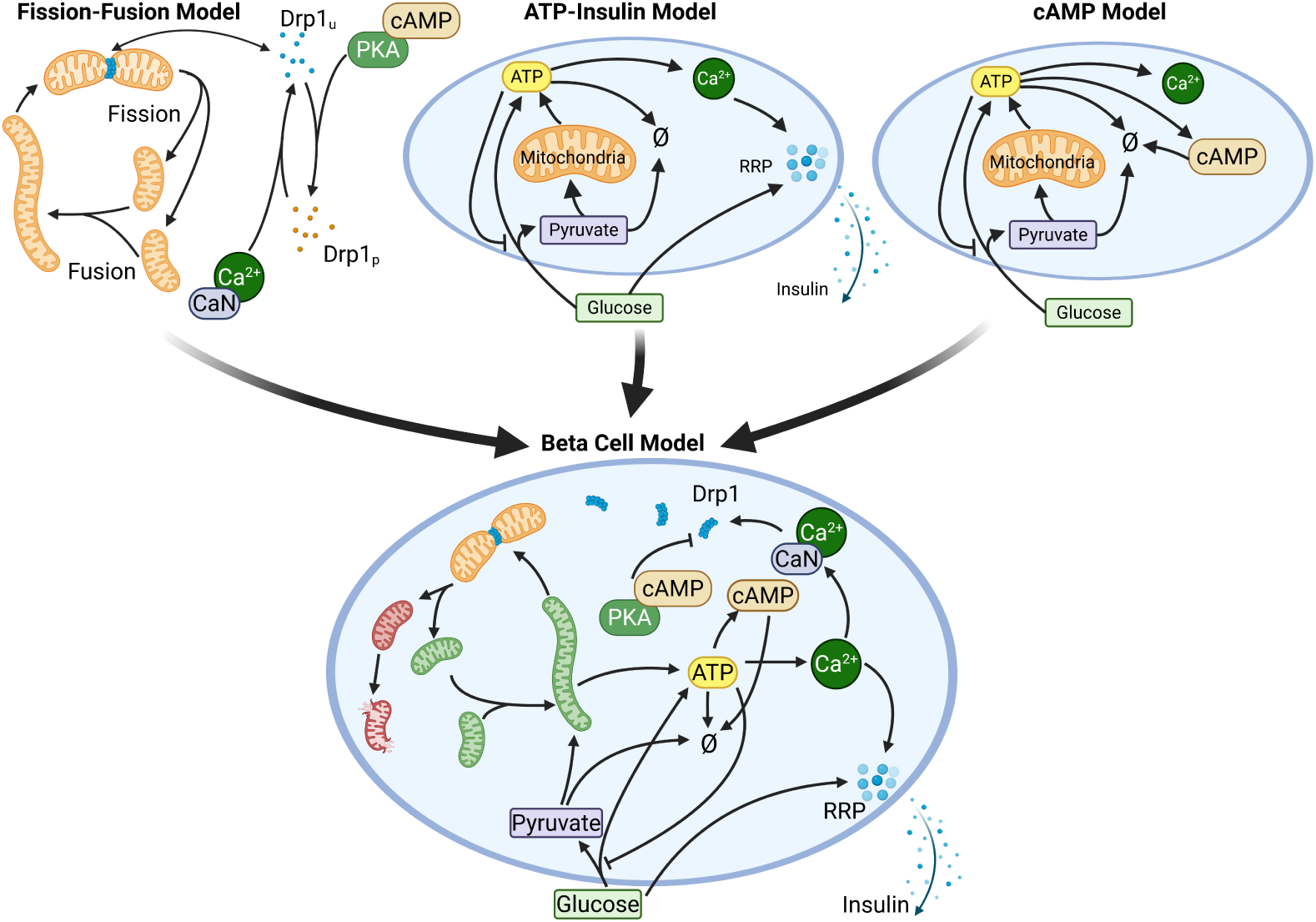
Sketch of the three submodels (fission-fusion model, ATP-Insulin secretion model, and cAMP model) and the beta-cell model. Created in BioRender. Henning, P. (2025) https://BioRender.com/2kh09q2.

#### ATP-Insulin secretion Model

This submodel (see Fig. 2, top center) includes the metabolism, Ca^2+^ flux, and insulin secretion of the beta cell. The model-building process is split into three phases. First, the ATP generation was modeled and calibrated. Here, glucose is metabolized to pyruvate and ATP via glycolysis [6, 7]. Pyruvate is either removed from the system to model anaerobe fermentation or used by the mitochondria to generate ATP via oxPhos [8–10]. The parameters that could not be taken from the literature were estimated based on data of the ATP content [40] and ATP production [41, 42] for different glucose levels in Min6 cells. In the second step, the calcium dynamic is added. A rise in the ATP level leads to an increase in the Ca^2+^ level. The response curve for this reaction could be taken from the literature [43]. Lastly, the insulin secretion dynamic is added and fitted. The increase in the calcium level triggers the insulin release from a rapid-release pool (RRP) [4]. The replenishment of the RRP is modeled as a glucose-dependent process. To fit the parameters, data on insulin secretion rates for different glucose levels in Min6 cells [44]were used. The fission and fusion dynamics of the mitochondria are not considered in the submodel.

#### cAMP Model

The third submodel (see Fig. 2, top right) is the simplest of the three and adds the synthesis and degradation of cAMP. It is based on the ATP generation and calcium flux from the ATP-Insulin submodel (Sec. 3.1). To this base, rules for the ATP-to-cAMP conversion [45, 46] and cAMP degradation [47, 48] have been added. The parameter estimation uses data of the cAMP content for different glucose levels in Min6 cells [49]. As in the ATP-Insulin secretion model, mitochondrial fission and fusion are not considered.

### 3.2 Beta-Cell Model

The complete beta-cell model integrates all components from the three submodels (see Fig. 2, bottom) with some modifications and additions. Some rules are extended, and new rules are added to capture the behavior that relates to the different submodels.

Among the rules that can be reused without changes are the rules regarding the (de)phosphorylation of Drp1, the calcium flux, the insulin signaling pathway, and the cAMP dynamics. The major extension of the beta cell model is the introduction of mitochondrial health and damage attributes to simulate the dynamics of their physiological state. These attributes abstractly represent a range of damage markers, such as reduced mitochondrial membrane potential, mtDNA lesions, oxidized proteins, and elevated reactive oxygen species (ROS) levels. Mitochondrial damage is introduced via ROS, a byproduct of the oxidative phosphorylation (oxPhos) [50]. In the beta cell model, this dynamic is represented by a rule that converts health into damage in proportion to the oxPhos rate. Damage is either repaired by the turnover of protein [51–53] or exchange of mtDNA [51, 54] or accumulates until the mitochondrion is removed via mitophagy [39]. To compensate for the loss of mitochondrial mass that is caused by the removal of a mitochondrion, we added the capability for mitochondria to grow. Both the rules for mitophagy and biogenesis are added to the beta cell model; neither exists in any of the submodels, as they relate to fission-fusion dynamics and ATP generation.

The rate of oxidative phosphorylation in each mitochondrion is extended by a health-to-damage-ratio dependent function *f* to account for the observation that damaged mitochondria are less active [54, 55]. We model *f* as a sigmoidal function:

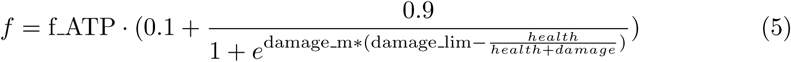

The parameter damage lim determines at which damage-to-health ratio the oxPhos rate decreases, and damage m determines how steep the decrease is. f ATP is a scaling factor compensating for ATP production losses introduced by coupling oxPhos to mitochondrial health.

For a detailed analysis, the assumption of one general anchor protein used in the fission-fusion submodel is replaced by the two known key factors, namely Fis1 and MFF. While MFF sites evoke midzone fission events, Fis1 sites trigger peripheral fission events. So the rule for fission now takes the specificity of the anchor protein into account. In addition, the peripheral fission mechanism is modified in this model such that the smaller daughter mitochondrion inherits a disproportionately higher level of damage, consistent with experimental observations [38]. For example, a quality asymmetry of 5% in a peripheral fission event that splits a mother mitochondrion at the ratio 25/75 would lead to a small daughter mitochondrion that carries 25%+5%=30% of the damage from the mother mitochondrion despite only having 25% of its volume. Due to the rule-based modeling approach of ML-Rules, this new behavior can be easily added. The peripheral fission rule from the fission-fusion submodel (without the kinetics, please see the appendix 4.3):

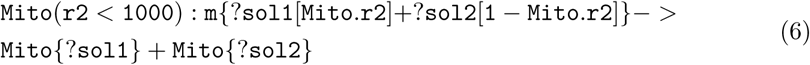

is changed to:

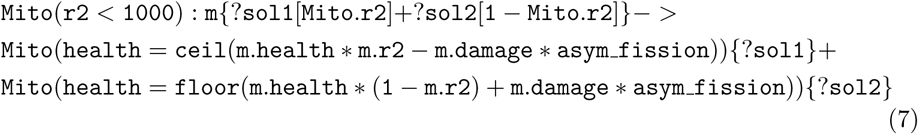

Here, the health attribute of the daughter mitochondria depends on the health of the mother mitochondrion (m.health), the fission position (r2), and (asym fission), which leads to an asymmetrical distribution of the health. The complete rule, also including the assignment of the damage and volume attribute, can be found in the appendix 4.3.

Finally, for the complete beta cell model, it is necessary to test whether the entire beta cell model still reproduces the data with which the original submodels were calibrated, thereby contributing to a valid composition [56]. As shown in the appendix 4.3, the complete beta cell model remains capable of reproducing the behavior of the submodels.

To assess the validity (or the range of application for which a model is valid), thorough documentation of the different sources used to generate the simulation model is important [57]. In our study, we used a provenance-based approach for documentation. The literature used to identify the key players and reactions, parameter values, and data for parameter estimation are shown in the provenance graph in Fig 3 which uses the provenance standard PROV-DM [58]. In the graph, the nodes represent entities (ovals), including references (yellow), models (blue), parameter estimation methods (green), and figures (pink), as well as activities (white rectangles), such as model refinement, calibration, or figure generation. The edges from the activities to the entities read as the activity uses this entity; the edges from entities to activities read as these entities are generated by this activity. E.g., the cAMP submodel (M3) is based on the ATP and calcium dynamics of the ATP-Insulin submodel (M2). A provenance graph also provides a good overview of the knowledge and data sources used to build models (and their interrelationships) [59].

**Fig 3.**
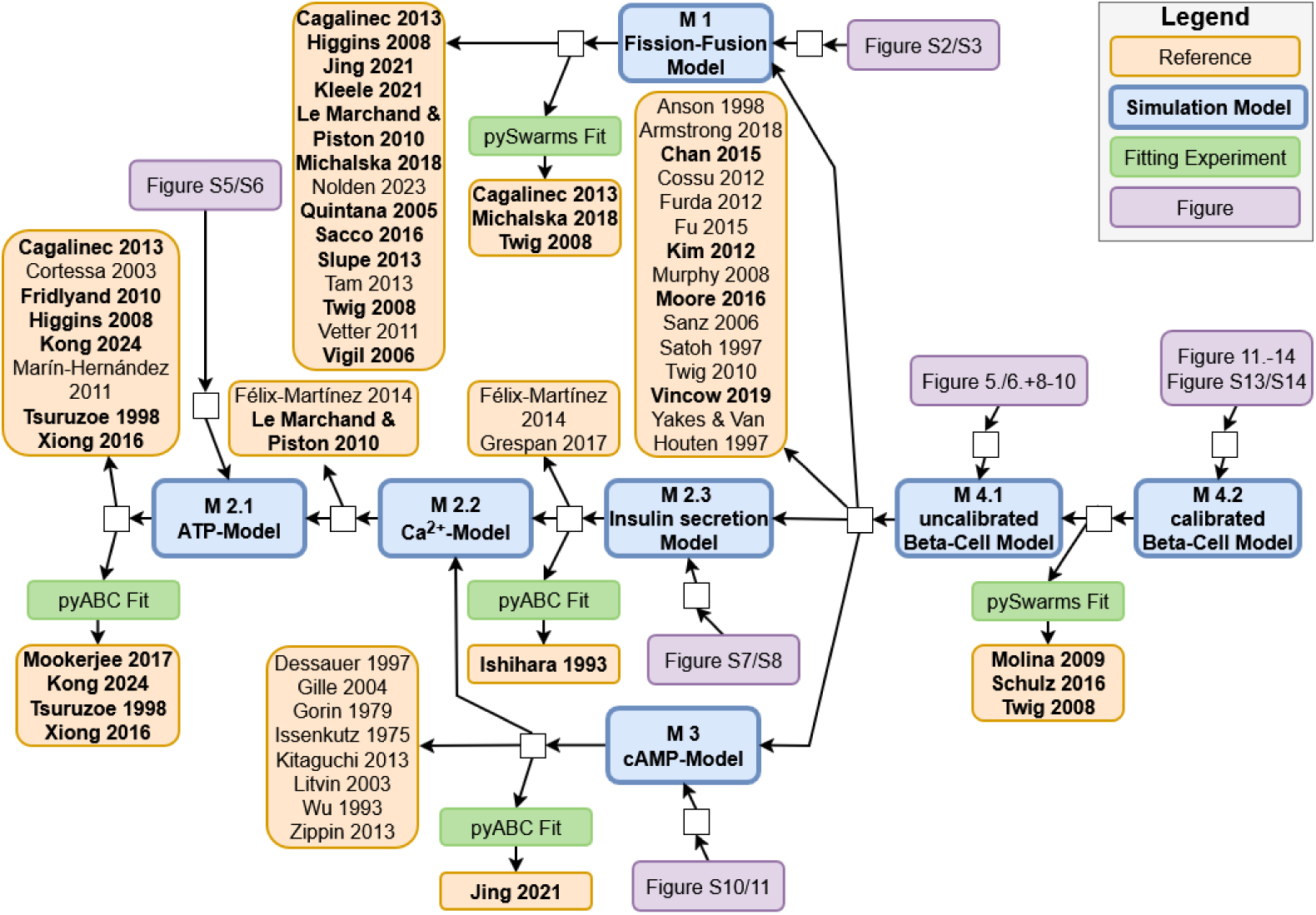
Provenance graph that shows the process of building the three submodels (M1-M3), the complete beta-cell model (M4), and the references used. From references written in bold, data such as concentrations or parameters were extracted.

## 4 Results

### 4.1 Damage Regulation by Damage-Sorting Fission

Our detailed beta-cell model allows us to investigate how fission, fusion, and mitophagy interact to regulate damage within the mitochondrial network. For effective quality control, one or more of these processes must exhibit selectivity with respect to mitochondrial damage. Mitophagy selectively removes small mitochondria that fit into an autophagosome [60] and have low membrane potential, which indicates respiratory chain damage [39].

For fusion events, it was observed that depolarized mitochondria could no longer fuse with other mitochondria [61] and, therefore, remain in a state where they can be potentially removed by mitophagy. Fission events can create daughter mitochondria with different membrane potential [39].

Previous simulation studies [12, 62] have investigated the underlying mechanism. In these simulation models, mitochondria exist in two states: solitary or fused, and carry either 10 or 20 health units, respectively. Each health unit can be either healthy or damaged. Before a fused mitochondrion is divided into two solitary ones, a fixed number of health units is exchanged randomly. This exchange can create solitary mitochondria that carry a large number of damaged health units. If these are removed from the network via mitophagy, the overall health of the network is improved. After the removal of a solitary mitochondrion, one of the remaining mitochondria is copied to keep the number of mitochondria constant during a simulation run.

In our model, in contrast to previous ones, the fission dynamics implement the more recent findings by Kleele et al. [38], which show that the amount of damaged parts is disproportionately high in small daughter mitochondria after a peripheral fission event (see Sec. 3.2 and Fig. 4). To examine the impact of this apparent sorting mechanism, we introduced two parameters. One parameter is the ”damage threshold”, at which mitochondria can no longer fuse with others and can be removed. The second parameter is the ”fission asymmetry”, which is short for fission damage asymmetry and determines how much damage is sorted into the small mitochondria during a peripheral fission event. We conduct a full factorial parameter scan over these two factors, with N = 5 replications per parameter configuration. Figure 5 presents the steady-state damage levels of the mitochondrial network for different values of the damage threshold and fission asymmetry. In the absence of a sorting mechanism during peripheral fission (Fig. 5: a) fission asymmetry = 0%, b) A=0%), the damage level in the network is roughly equal to the damage threshold (compare dark blue line and dashed line in Fig. 5 b)). An increase in the damage threshold always results in an increase in mitochondrial damage. As the fission asymmetry increases, more damage is concentrated in the smaller daughter mitochondria, which are removed via mitophagy, and the network-wide damage is decreased (Fig.5). A closer look into the damage levels of individual mitochondria, shown in Fig. 6, reveals that all mitochondria are close to the damage threshold in the absence of a damage sorting mechanism (blue bars). In this case, the mitochondria lack a mechanism to mitigate their damage. Both types of fission create daughter mitochondria with the same level of damage, and the removal of a small, damaged mitochondrion does not improve the overall damage level of the network. If the damage sorting mechanism is set to an asymmetry of 5% (orange bars) or 10% (yellow bars), the damage distribution is notably broader and shifted toward lower damage levels.

**Fig 4.**
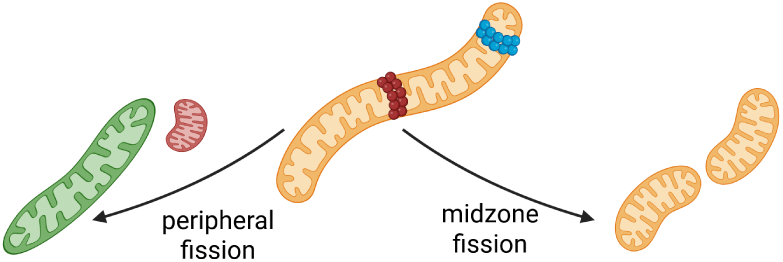
Schematic representation of the differences between a peripheral and a midzone fission event as they were observed by Kleele et al. [38]. Created in BioRender. Henning, P. (2025) https://BioRender.com/l0el4e4.

**Fig 5.**
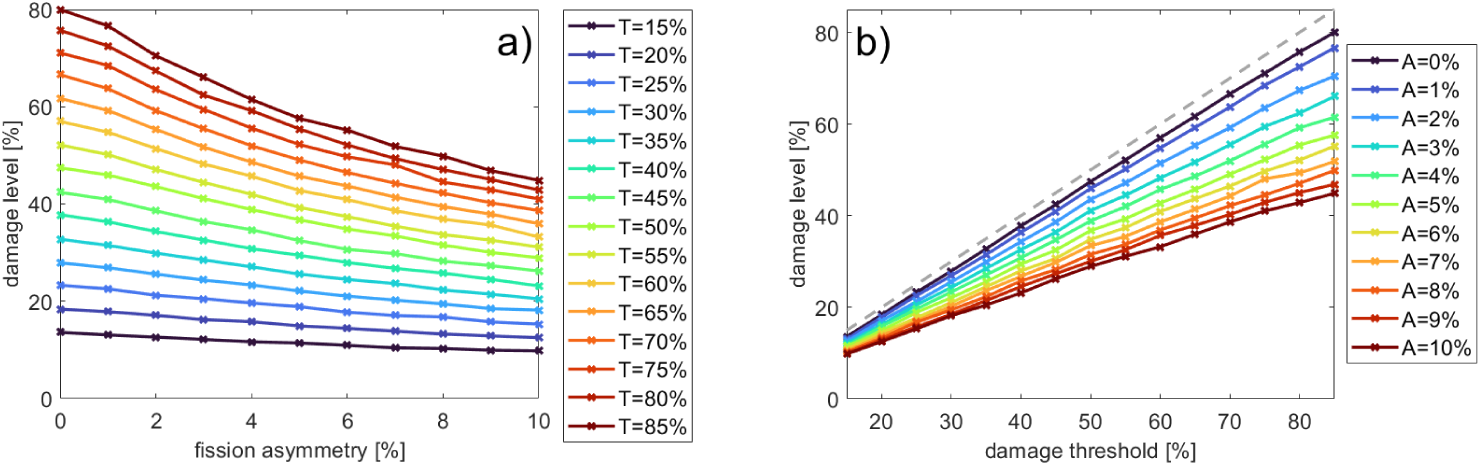
Steady-state damage levels in the mitochondrial network as a function of fission asymmetry (A) and damage threshold (T), averaged over N = 5 replicates. The grey dashed line in b) indicates the case where the steady-state damage of the network is equal to the damage threshold.

**Fig 6.**
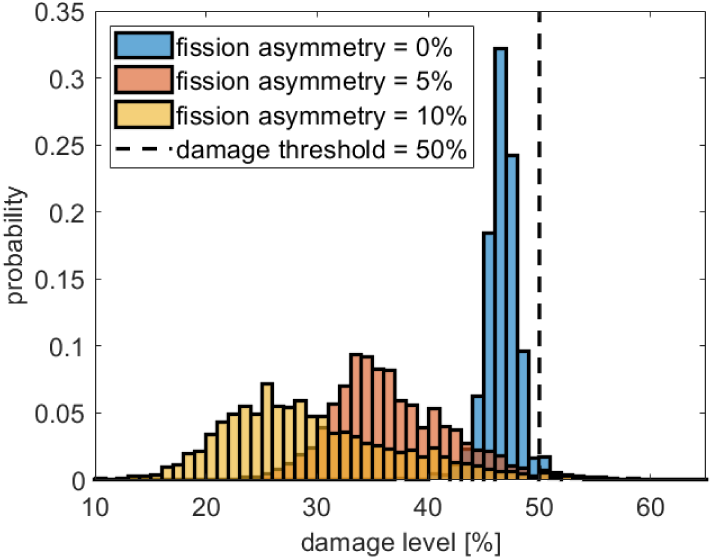
Distribution of the damage levels of individual mitochondria for the absence of damage sorting fission (fission asymmetry = 0%) and two different sorting mechanisms. The damage threshold was set at 50% for all simulations. N=5 replications.

In this case, a peripheral fission event of a mitochondrion close to the damage threshold might create a large sister mitochondrion with a notably reduced damage level and a small daughter mitochondrion with a damage level above the damage threshold that can be removed from the network.

An interesting feature of the damage sorting peripheral fission revealed by our simulation model is the observation that small changes in the fission asymmetry result in larger changes regarding the damage level. An increase of the fission asymmetry from 0% to 5% (at a damage threshold of 50%) decreases the damage level of the healthiest mitochondria in Fig. 6 from ∼40% to ∼25% and the average damage in Fig. 5 a) from

∼42% to ∼32%. This difference in the damage level can be attributed to the occurrence of successive, damage-sorting fission events that lead to an accumulated improvement, exceeding the effect of a single damage-sorting fission event.

### 4.2 Downregulation of Fis1 and MFF affects mitochondrial structure and damage levels differently

Having demonstrated that peripheral fission events with damage sorting are crucial for maintaining mitochondrial network health, we next investigated how the downregulation of the fission proteins Fis1 and MFF affects mitochondrial damage levels. As shown in Fig 7, Fis1 regulates peripheral fission events. The positive effect on the health state of the mitochondrial network was discussed in the previous section. MFF, on the other hand, creates two roughly equally sized mitochondria with a similar damage level. To determine the effect of both anchor proteins on mitochondrial damage, the system was simulated with a damage threshold of 50%, a fission asymmetry of 5% (so the smaller mitochondria resulting from a fission event carry 5% more damage), the amount of Fis1 and MFF was successively reduced from 320 to 0 (in steps of 20) while the other anchor protein was kept constant at 306 [21, 22]. For each configuration, five simulations were executed. Fig. 8 a) shows that the number of peripheral and midzone fission events decreases nearly linearly with the amount of Fis1 and MFF in the cell, respectively. A slight increase in the other fission type was observed in both cases, likely due to an increased pool of unbound Drp1.

**Fig 7.**
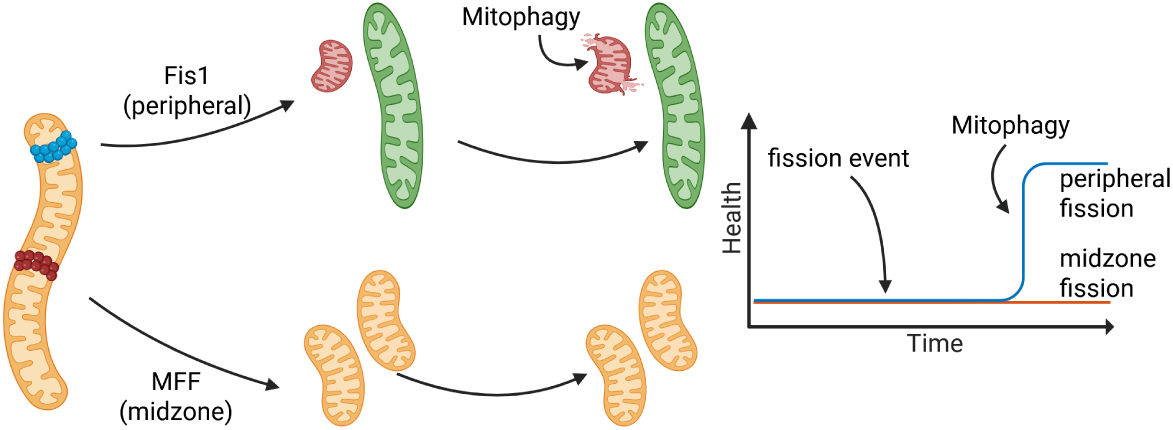
Sketch of peripheral and midzone fission event regulated by Fis1 and MFF, respectively. Fission events themselves do not change the health state of the mitochondrial network. However, the removal of the small damaged daughter mitochondrion, by mitophagy, enabled by a peripheral fission event (assuming damage sorting), would improve the health of the network. Created in BioRender. Henning, P. (2025) https://BioRender.com/5xl7gej.

**Fig 8.**
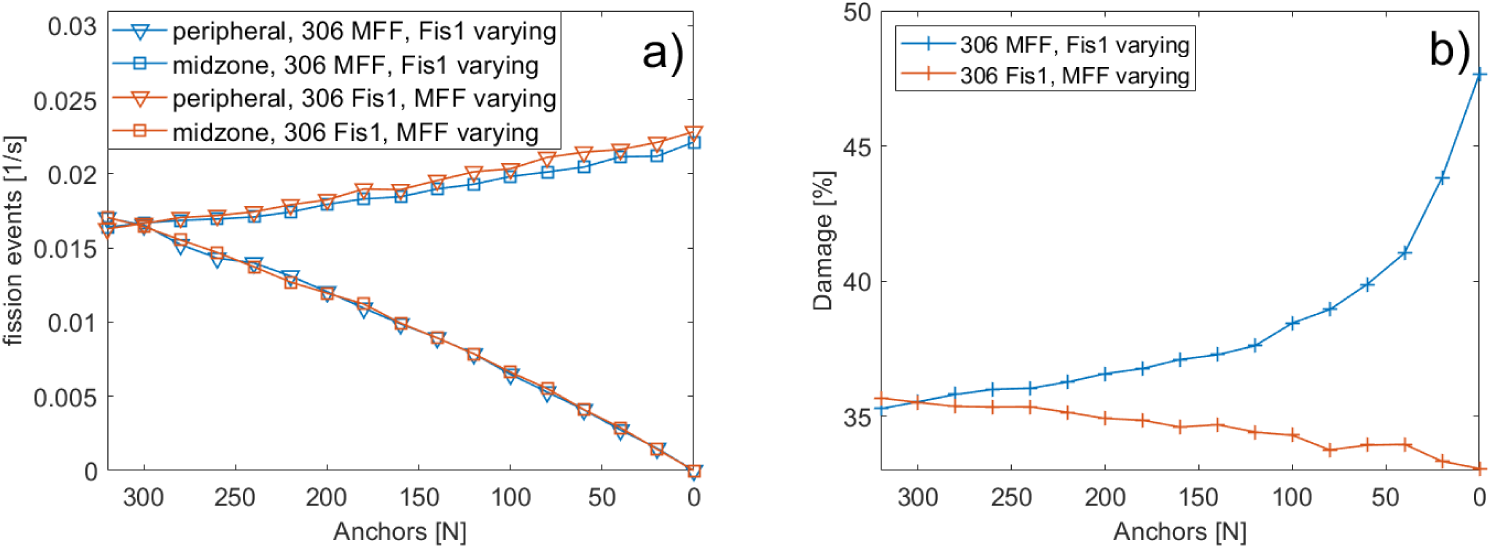
a) Fission events per second during Fis1 (blue) or MFF (orange) downregulation. The other anchor protein was fixed at 306. Triangles (∇) denote peripheral fission events; squares (□) denote midzone fission events. b) Steady-state damage level for the downregulation of Fis1 (blue) or MFF (orange).

The steady-state mitochondrial damage increased progressively as Fis1 levels declined (Fig. 8b), with a marked rise when Fis1 dropped below 40. In the case of zero Fis1 anchor sites and, consequently, no peripheral fission events, the damage level of the network is close to the damage threshold, as was also observed in Fig. 5 for the absence of a sorting mechanism in peripheral fission events. In contrast, MFF downregulation caused a slight reduction in overall damage, potentially due to the modest increase in peripheral fission, underlining again the importance of peripheral fission for quality control [38, 39].

A comparison of the distribution of individual mitochondrial damage for the cases of 0 or 300 Fis1 (Fig. 9 a)) shows that the downregulation of Fis1 leads to a shift of the damage distribution towards the damage threshold, similar to Fig. 6 and the absence of a sorting mechanism. On the other hand, the downregulation of MFF changes the distribution of mitochondrial damage to slightly lower levels(Fig. 9 b)).

**Fig 9.**
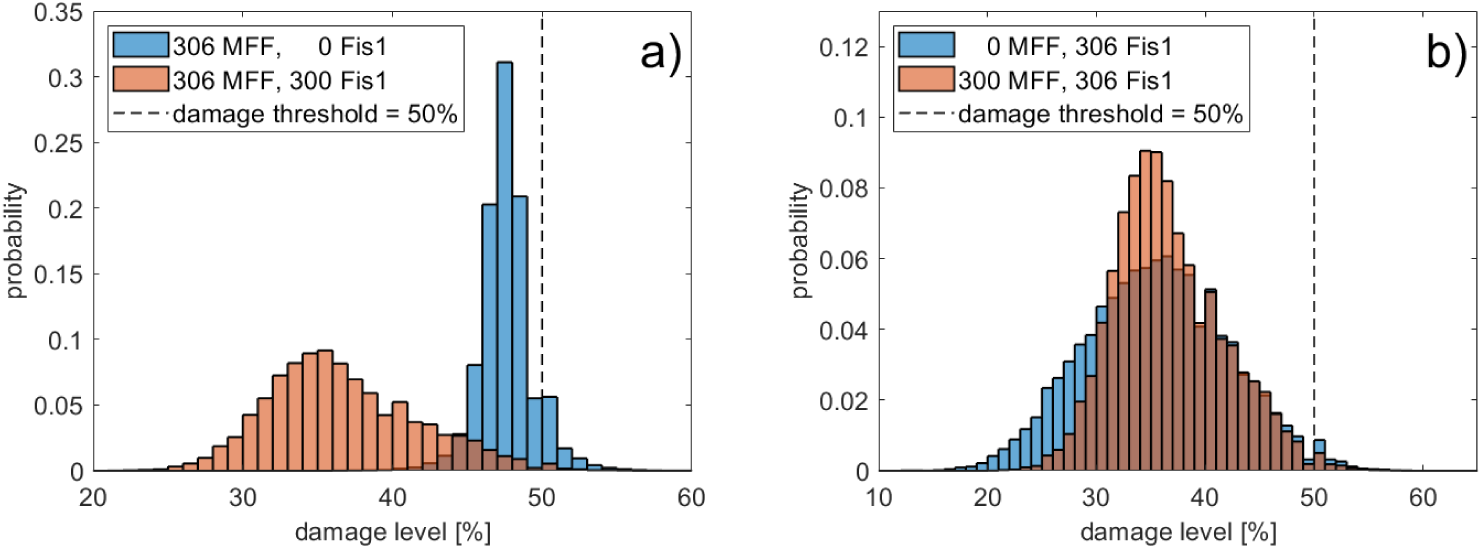
Distribution of the damage levels of individual mitochondria for the knock-out of Fis1 or MFF. Orange bars show the case of roughly equal amounts of MFF and Fis1. Blue bars show the knock-out of Fis1 a) or MFF b). The damage distribution was taken from the same runs shown in Fig. 8.

Finally, we examined the size distribution of mitochondria in the Fis1/MFF knockout case to determine how the two different fission types alter the structure of the mitochondrial network. The knockout of MFF and, therefore, the absence of midzone fission events (Fig.10, orange bars) leads to larger mitochondrial networks as this slightly promotes peripheral fission (see Fig. 8 a)). In this case, small and damaged mitochondria, created by peripheral fission, are removed from the network, while the larger and healthier part of the mitochondria remains and can fuse to create even larger ones. These findings align with the observation that MFF/midzone fission is associated with cell proliferation, which needs a more fragmented mitochondrial network to distribute the mitochondria evenly between the daughter cells [38, 63]. The knockout of Fis1 and, therefore, the absence of peripheral fission events leads to fewer large networks (Fig.10, blue bars) - a situation similar to that depicted in our motivation image (Fig. 1).

**Fig 10.**
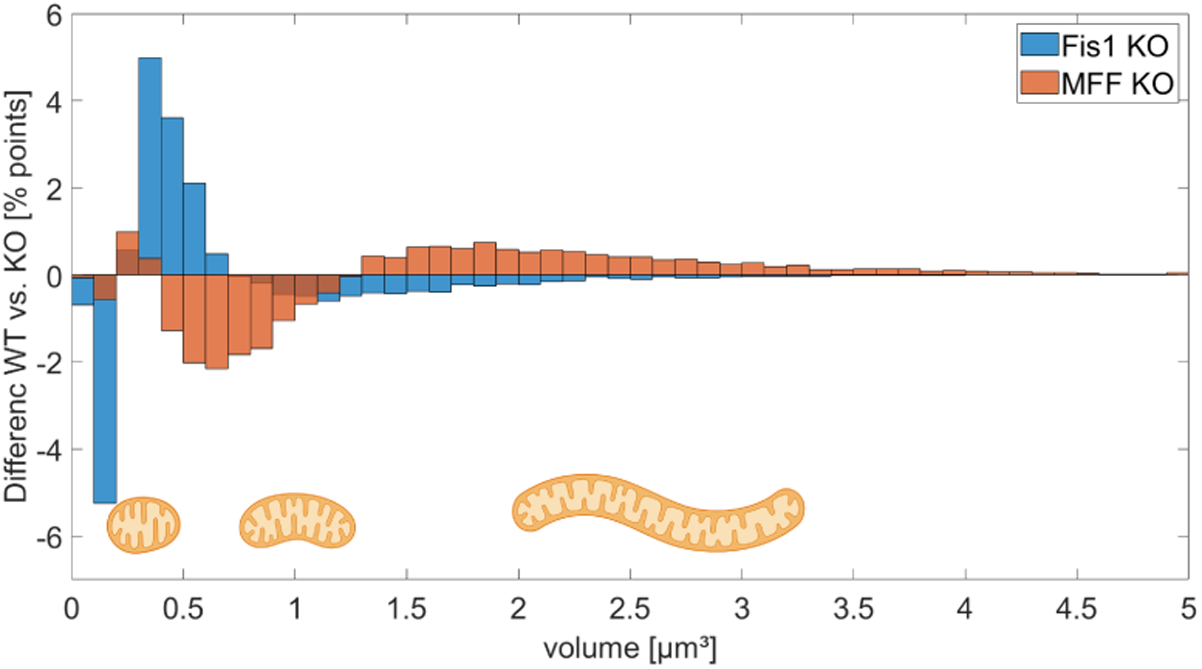
Changes in the distribution of the volume of the mitochondria between WT and Fis1 KO cells (blue) or WT and MFF KO cells (orange).Created in BioRender. Henning, P. (2025) https://BioRender.com/sestsgw.

This confirms that precisely regulated Fis1 expression is essential for maintaining a healthy mitochondrial network. In vitro studies have shown that the down-regulation of Fis1 decreases the health in the mitochondrial network [38] and the glucose-stimulated insulin secretion [39]. Furthermore, increasing the expression of Fis1 in glucose-unresponsive beta cells restores their functionality [64].

### 4.3 Interplay of Mitochondrial Damage and Insulin Secretion

After investigating the role of peripheral fission in regulating mitochondrial health and how this is affected by the downregulation of Fis1, we next aim to couple this effect to the insulin secretion of pancreatic beta cells. As discussed in section 3.2 and shown in several studies, significantly elevated ROS levels cause mitochondrial damage and a reduced oxygen consumption rate (OCR), and consequently a decreased ATP level [54, 65–69]. The rise of the ATP level is a crucial step in the insulin secretion pathway. Our model includes five parameters related to mitochondrial health and ATP synthesis capacity:

- f ATP: correction factor for ATP production
- damage lim: damage limit for the decrease of the oxPhos rate
- damage m: steepness of decrease of the oxPhos rate with damage
- asym fission: damage asymmetry during peripheral fission
- Thr damage: damage threshold at which mitochondria can no longer fusion and can undergo mitophagy

We first performed a global sensitivity analysis using Sobol’ indices to assess how these parameters influence insulin secretion. As shown in Fig. 11, damage lim, Thr damage, and asym fission exhibit the highest sensitivity. While damage lim and Thr damage show a first and second-order sensitivity, asym fission only shows a second-order effect, particularly in combination with Thr damage. These results are plausible as damage lim is directly coupled to the production of ATP and thus to the secretion of insulin. The other two parameters are vital for the health of mitochondria, as shown before, which is again essential for an effective production of ATP.

**Fig 11.**
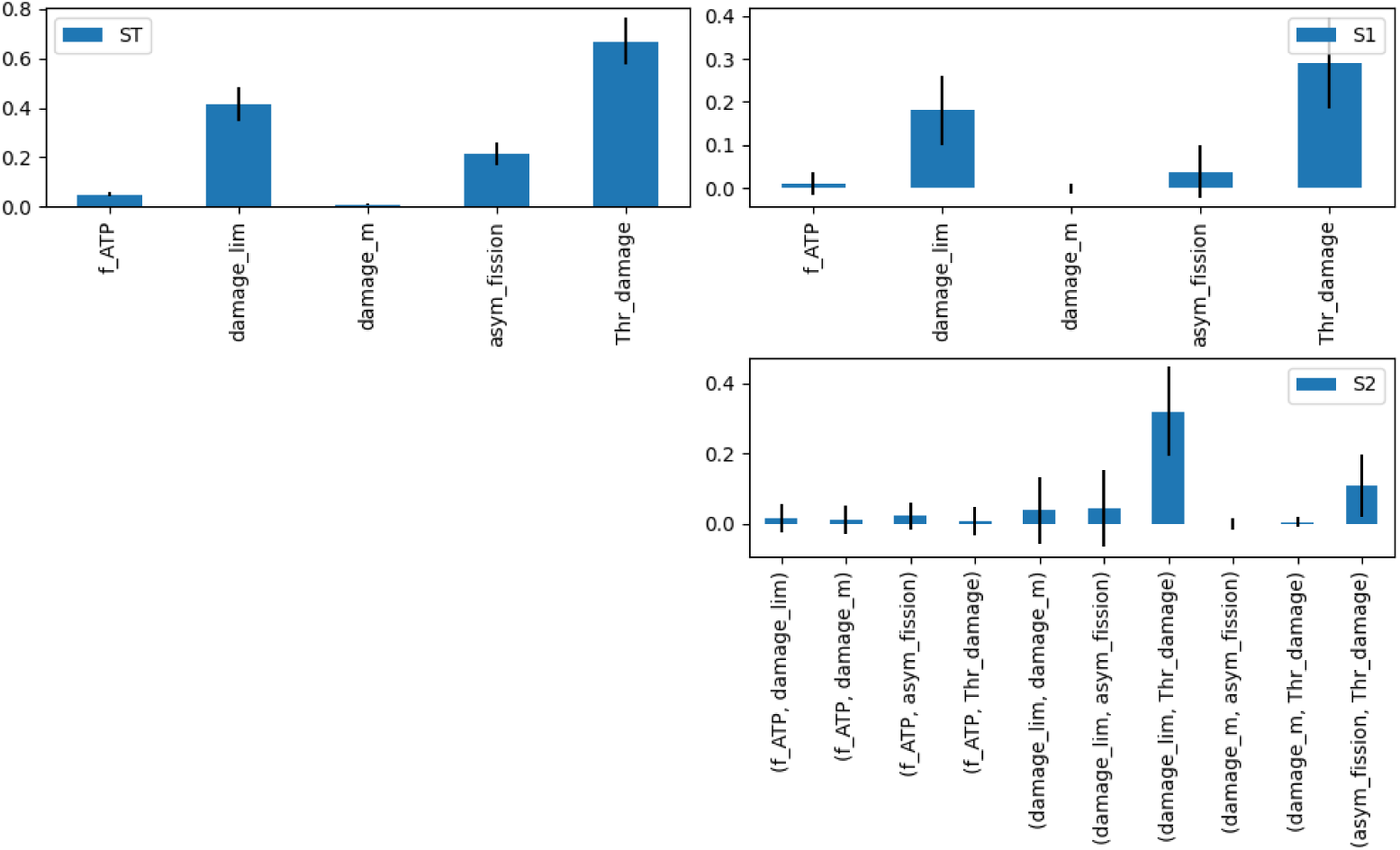
Sobol’ indices for the total sensitivity (ST), first-order sensitivity (S1), and second-order sensitivity (S2) of the five parameters f ATP, damage lim, damage m, asym fission, and Thr damage concerning insulin secretion.

Next, we aimed to determine which configuration of these five parameters best reproduces the insulin secretion observed in Fis1 knockdown (KD) cells. Experimental findings show a reduction in the response [39, 70] up to the beta-cell being non-responsive to high glucose levels [64]. In all these studies, basal insulin secretion remains unaffected. For our fitting experiments, we aim for a 40% reduction in insulin secretion at 25 mM glucose compared to wild-type cells [44].

The best-fit parameters are shown in Table 1. From the three parameters that affect the insulin secretion, the value of asym fission is close to the upper boundary set for the parameter fitting. We decided against an increase of the boundary, since this could lead to the case where all small mitochondria created are highly damaged. However, it is reported that some of the small mitochondria show normal membrane potentials and ROS levels [38]. Figure 12 compares the insulin secretion at different glucose concentrations for WT and Fis1-KD conditions. Both models exhibit a similar insulin secretion at low glucose levels (up to 5 mM) as reported in the literature [39, 64, 70]. At higher glucose concentrations, the WT model closely matches the experimental data. The Fis1-KD model shows a ≈34% reduction in secretion at 25 mM, which is also consistent with reported knockdown effects [39, 70] and a delayed response at ≈10 mM glucose. For the same set of parameters, we investigated how knocking down Drp1 would affect insulin secretion. As shown in Fig. 13, fission frequency declines steeply with Drp1 downregulation, more rapidly than with Fis1 or MFF (cf. Fig.8a). Furthermore, the fission rate drops to zero before the number of Drp1 molecules does. At these low levels of Drp1, it can no longer polymerize into rings of sufficient size to trigger a fission event. The mitochondrial damage increases with Drp1 depletion, reaching near-total damage at 5000 Drp1_10_ units (see Fig. 14a)). In contrast, damage levels for the entire network above the damage threshold were not observed in the knockdown of Fis1 or MFF (see Fig. 8b). Recall that the damage threshold determines which mitochondria can be subject to degradation. In the case of Fis1 knockdown, even the remaining fission activity of MFF (midzone fission without a damage sorting mechanism) was enough to keep the damage level slightly below the damage threshold. Accordingly with Drp1 depletion, the insulin secretion shown in Fig. 14b)) decreases more steeply than the damage rises and plateaus at a very low level, which corresponds to the basal level. The results of Drp1 knockdown studies on the insulin secretion vary. Reinhardt et al. [71] observed complete loss of glucose responsiveness in shDrp1 INS1 cells, while Kabra et al. [72] reported attenuated but present glucose response in shDrp1 MIN6 cells. The basal insulin secretion at low glucose levels remains unaffected in both publications. Our model aligns with the former, replicating the severe impairment of glucose-stimulated insulin secretion at a reduction of Drp1 of ≈25%.

**Fig 12.**
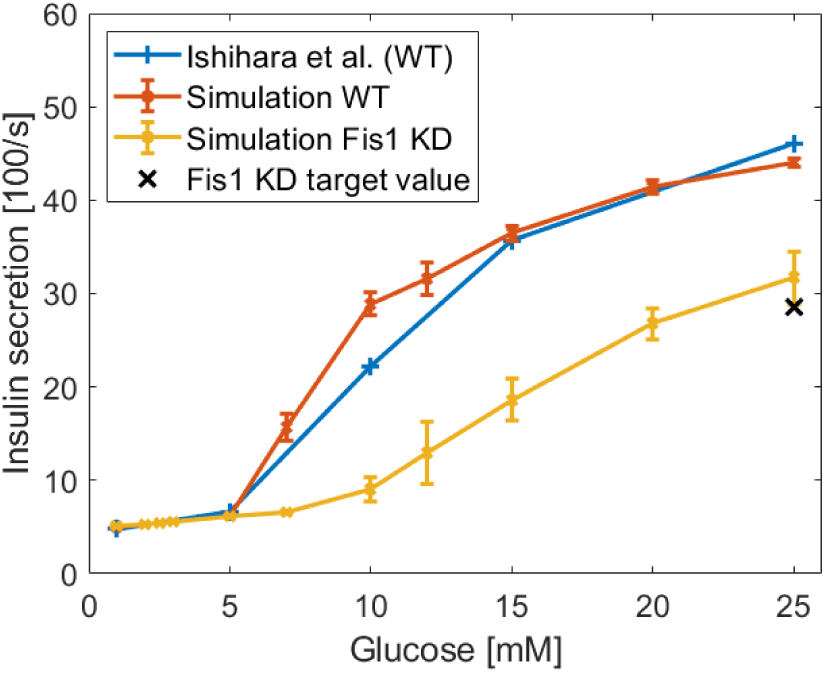
Simulated insulin secretion at different glucose concentrations for WT (orange) and Fis1 KD (yellow) cells, compared to wet-lab data (blue) [44]. The black cross represents the target value for the Fis1 knockdown case, with a 40% reduction in insulin secretion.

**Fig 13.**
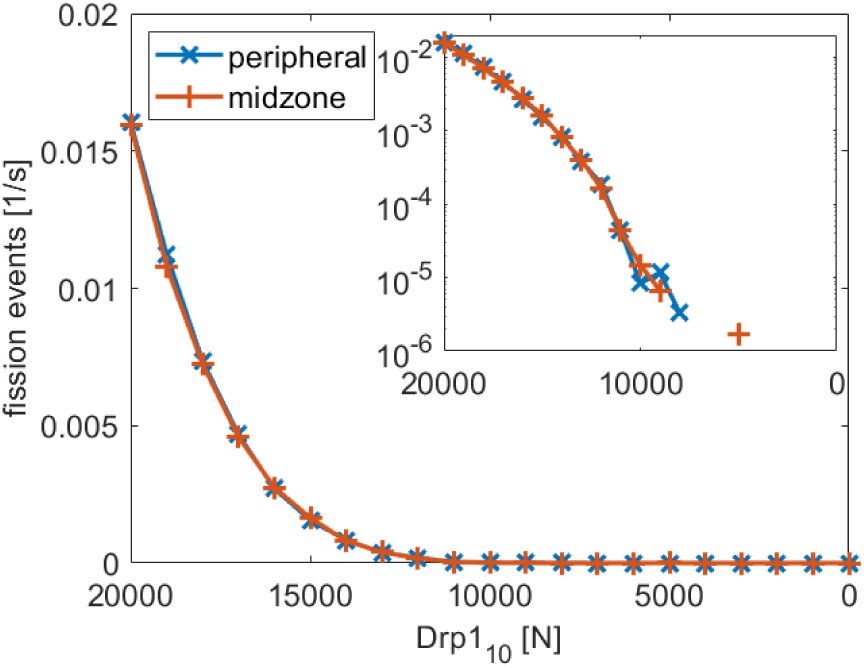
Peripheral (blue) and midzone (orange) fission rates as a function of Drp1_10_ clusters. 20000 Drp1_10_ corresponds to the WT case shown in Fig. 12. The insert shows the same data on a logarithmic y-scale. Missing values in the insert indicate zero fission events.

**Fig 14.**
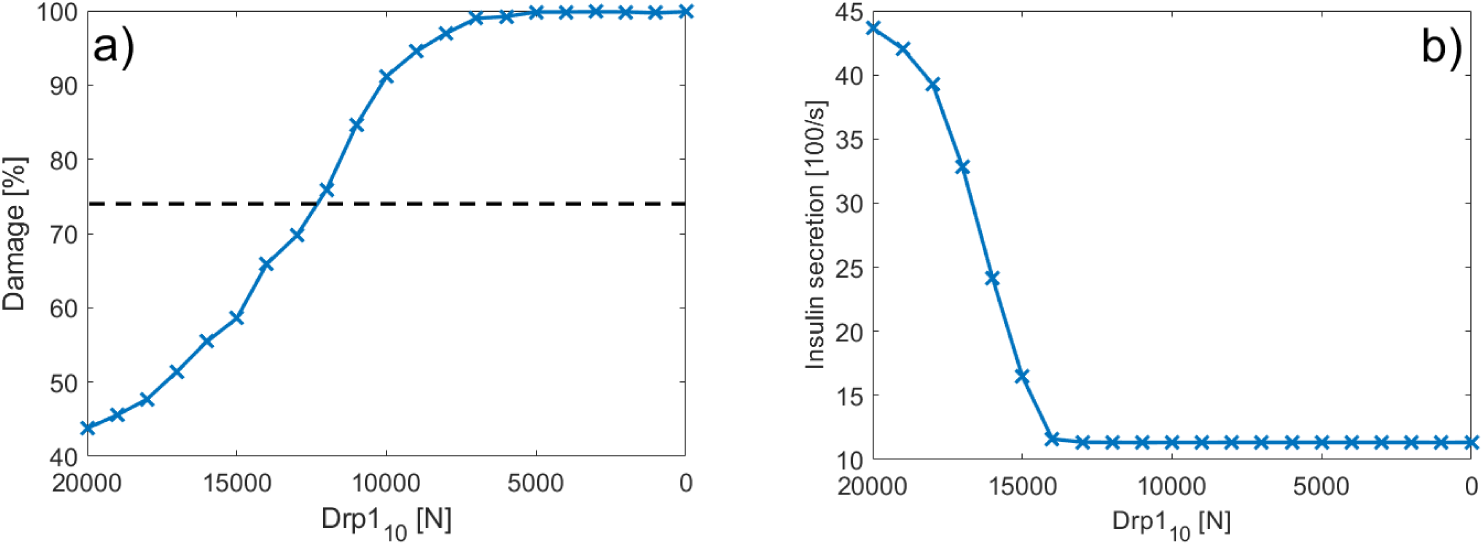
a) Mitochondrial damage as a function of Drp1_10_ concentration. The dashed line shows the damage threshold of 74% (see Tab. 1) b) Insulin secretion at [Gluc] = 25 mM for different Drp1_10_ levels. In a) and b) the case of 20000 Drp1_10_ corresponds to the WT case shown in Fig. 12

**Table 1.** Fitted parameter values to model the effect of a Fis1 knockdown on insulin secretion.

| parameter | value | boundaries |
| --- | --- | --- |
| f_ATP | 1.54 | 0.9 - 1.6 |
| damage_lim | 53.3% | 25% - 75% |
| damage_m | 28.7 | 10 - 50 |
| asym_fission | 9.22% | 0% - 10% |
| Thr_damage | 74.0% | 10% - 90% |

Overall, maintaining a balanced expression of both Fis1 and Drp1 in pancreatic beta cells is crucial for proper cellular function. Some studies observe an upregulation of Drp1 in response to high levels of glucose and /or fatty acids, leading to a high level of mitochondrial fragmentation and eventually to mitochondrial damage [73–76]. Although our model aims at investigating the effects of Fis1 knockdown on mitochondrial dynamics and insulin secretion, the results on the downregulation of Drp1 in Fig. 14 are a first step into a detailed investigation of the role of varying Drp1 expressions in these processes with our model. Thus, it contributes to our understanding of balancing regulatory proteins, which is difficult to achieve through an experimental approach alone.

## Conclusion

We developed a multi-level simulation model of pancreatic beta-cells that integrates insulin secretion, intracellular and mitochondrial metabolism, and mitochondrial dynamics. Exploiting the multi-level rule-based modeling and simulation approach of ML-Rules enabled us to integrate all these dynamics within a single model using one formalism. Unlike previous models, our model explicitly incorporates the fission proteins Fis1, MFF, and Drp1, enabling mechanistic insights into their distinct roles. Building on experimental findings, we differentiate between peripheral fission (Fis1-regulated, damage-control) and midzone fission (MFF-regulated, proliferation-related).

Our model focuses on damage regulation via fission, excluding the role of mitochondrial fission during cell proliferation. We investigate how damage sorting during peripheral fission affects mitochondrial functionality. We show that directing damaged components into small mitochondria is essential for maintaining mitochondrial health. Simulations also reveal how knockdowns of Fis1, MFF, and Drp1 alter mitochondrial health and morphology. MFF downregulation mainly reduces midzone fission, slightly affects damage levels, and increases the number of large mitochondria. Fis1 downregulation disrupts peripheral fission, leading to increased damage and the more small mitochondria. Drp1 downregulation eliminates all fission events, driving damage to ≈ 100%. We linked mitochondrial health to ATP production and could show reduced insulin secretion after Fis1 knockdown, while Drp1 knockdown renders beta cells glucose-unresponsive due to severe mitochondrial damage.

Overall, our findings underscore the essential role of mitochondrial quality control, particularly through damage sorting and peripheral fission, in maintaining beta cell function. Explicitly modeling fission proteins provides a mechanistic framework for understanding how disrupted mitochondrial dynamics contribute to the development of metabolic diseases.

Within our model, different processes are represented at varying levels of detail. Fission is modeled comprehensively, whereas fusion is simplified, considering only mitochondrial health and volume. A more detailed fusion model incorporating OPA1 and MFN1/2 would enable the exploration of the fusion-fission interplay in regulating glucose metabolism and insulin signaling [2]. Mitophagy is also simplified, considering only mitochondrial size and health. Integrating the Pink1-Parkin pathway and lysosomal mechanisms would enable the study of mitophagy dynamics under Drp1 upregulation [76]. The model’s multi-level rule-based design supports such extensions and the integration of new data.

We thus present an approach that enhances our understanding of mitochondrial quality control in beta cells. It will help develop new therapeutic strategies to mitigate mitochondrial damage and restore insulin secretion in type 2 diabetes.

## Supporting information

Appending and model files.

## Supporting information

**S1 Appendix. Detailed description of the (sub)models, the parameters, and the parameter estimation.**

**S2 File. Model file of the fission-fusion submodel.**

**S3 File. Model file of the ATP-Insulin submodel.**

**S4 File. Model file of the cAMP submodel.**

**S5 File. Model file of the beta cell model.**

**S6 File. pySwarms script for the estimation of the parameters of the fission-fusion submodel.**

**S7 File. pyABC script for the estimation of the parameters of the ATP dynamics of the ATP-Insulin submodel.**

**S8 File. pyABC script for the estimation of the parameters of the Insulin secretion dynamics of the ATP-Insulin submodel.**

**S9 File. pyABC script for the estimation of the parameters of the cAMP submodel.**

**S10 File. pySwarms script for the estimation of the parameters of the complete beta cell model.**

## Acknowledgments

The authors would like to thank Till Köster for implementing and optimizing ML-Rules3. Without this tool, the model presented here could not have been developed. We are grateful to Fiete Haack for his feedback on the manuscript. In addition, the authors are grateful to Mark Philipp and Clemens Schafmayer (Department of General, Visceral, Thoracic, Vascular and Transplant Surgery, University Medical Center Rostock, Rostock, Germany) for providing the human pancreatic tissue. SB and JS acknowledge funding from the German Diabetes Association (DDG). PH and AU acknowledge funding from the Deutsche Forschungsgemeinschaft (DFG, German Research Foundation, https://www.dfg.de/en) grant 225222086. In addition, PH received funding from the Young Neuro Scientist Programme of the Centre for Transdisciplinary Neurosciences Rostock (CTNR).

## Ethics statement

Human adult pancreatic islets were donated from biopsies performed during pancreatic surgery, as approved by the ethics committee of University Medicine Rostock (reference number 2019-0187).

