## Appending and model files. for "Illuminating the Role of Asymmetric Mitochondrial Fission on Beta-Cell Health": S1_appendix.pdf


### 1 Submodels

In this section, the different submodels are described in detail. The literature used to derive the rules, rates, and rate parameters is presented.

#### 1.1 Fission-Fusion Model

The fission-fusion submodel focuses on the dynamics of mitochondrial fission and fusion. It is used to estimate the parameters of the binding and unbinding of Drp1 to an anchor site, as well as the fission rate. Fig. 1 shows a schematic representation of the model. To reduce the computational cost of the simulation, Drp1 is clustered into bundles of 10 particles.

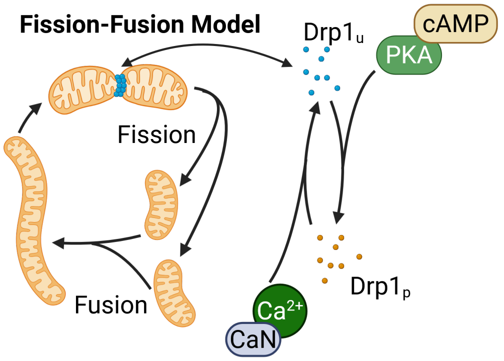


**Fig 1.** Schematic representation of the fission-fusion model. Unphosphorylated $Drp1_{10}$ clusters (blue dots) bind to and dissociate from anchor sites on the mitochondrial membrane. When a sufficient amount of Drp1_10 accumulates on an anchor site, a fission event can occur. Mitochondria can also undergo fusion, establishing a continuous fission-fusion cycle that constantly remodels the mitochondrial network. Drp1 is phosphorylated by PKA, which is activated by cAMP, and dephosphorylated by CaN, activated by $Ca^{2+}$. Created in BioRender. Henning, P. (2025) `https://BioRender.com/77bgd7a`

#### 1.1.1 Species

| Species | Description |
|---|---|
| Cell | beta-cell |
| Mito(rand1:float*, rand2:float*) | Mitochondrion, rand1/2: used for fission |
| Mito_vol | volume particle $\widehat{=}$ 0.01 $\mu m^3$ |
| Drp1_10(p:bool) | 10 Drp1 proteins, p: phosphorization state |
| Anchor | anchor site for Drp1 |
| PKA | cAMP-dependent protein kinase |
| cAMP | cyclic AMP |
| CaN | Calcineurin |
| Ca_100 | 100 $Ca^{2+}$ |

**Table 1.** Species used in the fission-fusion model.

#### 1.1.2 Rules

**Drp1 binding**
The Dynamin-related protein 1 (Drp1) is a key factor in the mitochondrial fission dynamics. It can oligomerize into rings around a mitochondrion, as long as it is unphosphorylated [1], and constrict to initiate a fission event. However, Drp1 alone cannot bind to the outer membrane of the mitochondria but needs an anchor protein. It was shown that binding of Drp1 to the anchor protein is not the limiting factor of mitochondrial fission, but the oligomerization of Drp1 [2]. Therefore, we only model the oligomerization step of Drp1 binding, assuming that prior binding to the anchor protein has already occurred and does not limit the reaction rate.

```
Mito{Anchor} + Drp1_10(p==false) -> Mito{Anchor{Drp1_10}} @ kon/vol;
```
(1)

**Drp1 unbinding**
Bound Drp1 can dissociate from anchor sites, following an isodesmic assembly model [3,4].

```
Mito{Anchor:a{Drp1_10}} -> Mito{Anchor} + Drp1_10 @@ 1/10*koff*#a;
```
(2)

**Drp1 phosphorylation**
The phosphorylation of Drp1 is catalyzed by the cAMP-dependent protein kinase (PKA), which is activated by cAMP [1]. Reaction kinetics follow Michaelis-Menten dynamics with a Hill-function-like activation by cAMP [5,6].

```
Cell{Drp1_10(p==false):d + PKA:p} -> Cell{Drp1_10(p=true) + PKA}
 @@ 1/10*kcatpka*#p*#d*10/(KMpka*vol*NA+#d*10)*100*##(cAMP_100 in Cell)
 /(K_act_PKA+100*##(cAMP_100 in Cell));
```
(3)

**Drp1 dephosphorylation**
Dephosphorylation of Drp1 is catalyzed by Calcineurin (CaN) [1], which is activated by $Ca^{2+}$. As for PKA, the kinetics follow Michaelis-Menton dynamics [7] with a Hill-function-like activation by $Ca^{2+}$ [8].

```
Cell{Drp1_10(p==true):d + CaN:c} -> Cell{Drp1_10(p=false) + CaN}
 @@ 1/10*kcatCaN*#c*#d*10/(KMCaN*vol*NA+#d*10)*100*##(Ca_100 in Cell)
 /(K_act_CaN+100*##(Ca_100 in Cell));
```
(4)

**Mitochondrial fission**

A fission event can occur once sufficient Drp1 has accumulated at an anchor site [3]. Mitochondria can undergo two different types of fission. The first is the midzone-fission event, where two roughly equal sister mitochondria are formed [9]. The fission is executed in two steps. First, the fission position is randomly generated (and saved in `rand1`) if a sufficient amount of Drp1 has accumulated in an anchor site and the Drp1 bound in this site returns to the cell compartment. The rate of the fission event depends on the size of the mitochondria, with larger ones being more likely to undergo fission [10, 11].

```
Mito:m{Anchor{Drp1_10:drp + ?solA}} ->
 Mito(rand1 = RandomGauss(0.5, 0.075)){Anchor} + Drp1_10+?solA      (5)
 @@ if (#drp>N_fission) then k_fission*(#(Mito_vol in m))^3;
```

Immediately after the position of the fission event is determined, the fission is executed, and the content of the mother mitochondrion is distributed between the two sister mitochondria based on the fission position rand1.

```
Mito(rand1>0):m{?sol1[Mito.rand1] + ?sol2[1-Mito.rand1]} ->
 Mito(rand1=-1, rand2=-1){?sol1} + Mito(rand1=-1, rand2=-1){?sol2}   (6)
 @@ immediately;
```

Equivalent rules govern the second type of fission events, peripheral fission. Here, the mitochondrion is divided into a small and a large sister mitochondrion. The position of the fission is stored in `rand2`.

```
Mito:m{Anchor{Drp1_10:drp + ?solA}} ->
 Mito(rand2 = RandomGauss(0.2, 0.04)){Anchor} + Drp1_10+?solA      (7)
 @@ if (#drp>N_fission) then k_fission*(#(Mito_vol in m))^3;
```

```
Mito(rand2>0):m{?sol1[Mito.rand2] + ?sol2[1-Mito.rand2]} ->
 Mito(rand1=-1, rand2=-1){?sol1} + Mito(rand1=-1, rand2=-1){?sol2}   (8)
 @@ immediately;
```

**Mitochondrial fusion**

The fusion of mitochondria is modeled as a simple rule with a constant reaction rate. In contrast to the fission rate, the fusion rate does not depend on the size of mitochondria [10].

```
Mito:m1{?solm1} + Mito:m2{?solm2} ->
 Mito(rand1=-1, rand2=-1){?solm1 + ?solm2} @@ k_fusion;              (9)
```

#### 1.1.3 Initial conditions

| Species | Number | Ref. |
|---|---|---|
| Cell | 1 | - |
| Mito(rand1 = −1, rand2 = −1) | 68 | [12] |
| Mito_vol | 100 per Mito | [12] |
| Drp1_10(p = true) | 10000 | [13] |
| Drp1_10(p = false) | 10000 | [13] |
| Anchor | 612 | [14, 15] |
| PKA | 300000 | [13] |
| cAMP | 10000 | [16] |
| CaN | 350000 | [13] |
| Ca_100 | 500 | [17] |

**Table 2.** Initialization of the fission-fusion submodel.

#### 1.1.4 Parameters

| Parameter | Description | Value | Ref. |
|---|---|---|---|
| kon | Binding rate or Drp1 | 0.025 $\mu m^3/s$ | fit to [3, 10, 12, 18] |
| Kd | Equilibrium constant for Drp1 binding | 0.301 $\mu mol/s$ | fit to [3, 10, 12, 18] |
| koff | Unbinding rate of Drp1 | kon ∗ Kd | - |
| k_fission | Rate coefficient for fission | $8.211{\cdot}10^{-8} Hz$ | fit to [3, 10, 12, 18] |
| N_fission | Number of Drp1 needed for fission | 11 | [3] |
| rand1 | Position of midzone fission | N(0.5, 0.075) | [9] |
| rand2 | Position of peripheral fission | N(0.2, 0.004) | [9] |
| k_fusion | Rate coefficient for fusion | $5.41{\cdot}10^{-4}$ 1/min | [10, 18] |
| vol | Volume of a Min6 cell | 1025 $\mu m^3$ | [19] |
| kcatCaN | Rate constant for CaN | 7.6 $1/s$ | [7] |
| KMCaN | Michaelis constant for CaN | 160 mM | [7] |
| K_act_CaN | Activation constant of CaN | 14 nM | [8] |
| kcatpka | Rate constant ok PKA | 4 1/s | [20] |
| KMpka | Michaelis constant for PKA | 70 mM | [20] |
| K_act_PKA | Activation constant of PKA | 0.9 $\mu M$ | [5] |
| NA | Avogadro constant | 6.022 1/mol | - |

**Table 3.** Parameters used in the fission-fusion submodel.

#### 1.1.5 Parameter Estimation

The submodel was used to estimate the rate coefficient for Drp1 binding (kon), the equilibrium constant of Drp1 binding (Kd), and the rate coefficient for mitochondrial fission (k_fission). PySwarms [21] was used to minimize the relative quadratic difference

$$error = \sum_i \left( \frac{sim_i - lab_i}{lab_i} \right)^2 \tag{10}$$

where $sim_i$ represents the simulated model output and $lab_i$ the corresponding experimental (wet-lab) data. The calibration targeted three experimental observables:

(i) the Drp1 assembly kinetics (2.6 $s^{-1}$) [3], (ii) the mitochondrial fission frequency (0.04 events $min^{-1}$ $mitochondrion^{-1}$) [10, 18], and (iii) the average number of mitochondria in a MIN6 cell (68 mitochondria) [12]. The resulting parameter values are listed in Table 3, together with parameters directly taken from the literature.

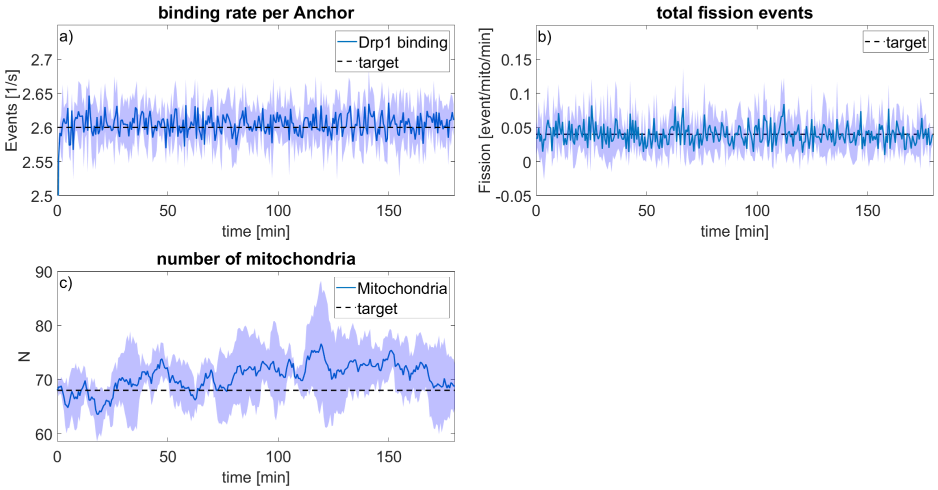


**Fig 2.** Comparison between literature data and simulation results for a) the Drp1 binding rate [3], b) the mitochondrial fission rate [10, 18], and c) the number of mitochondria in a Min6 cell [12]. The shaded area shows the standard derivation of N=5 simulation runs.

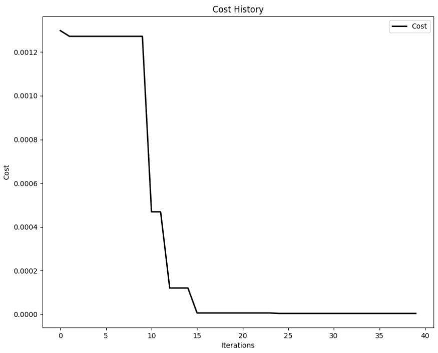


**Fig 3.** Cost function of the pySwarms run to estimate the parameters of the fission-fusion model.

The simulation results of the fission-fusion submodel are shown in Fig. 2. After the parameter estimation, the submodel successfully reproduces the experimental target values for the Drp1 binding rate, mitochondrial fission rate, and the number of mitochondria. Fig. 3 shows the evolution of the error function during optimization. After approximately 15 iterations, the error converged, and an optimal parameter set was obtained.

### 1.2 ATP-Insulin secretion model

The ATP-Insulin submodel focuses on ATP production and consumption, calcium signaling, and insulin secretion, as shown in Fig. 4. It is used to estimate parameters for the ATP dynamic and the insulin secretion dynamic.

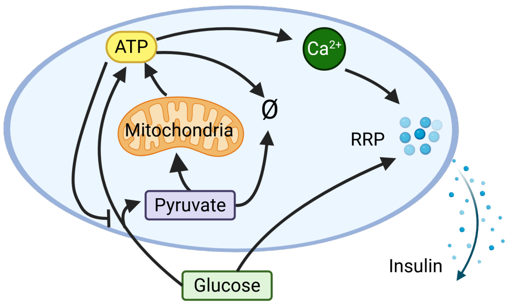


**Fig 4.** Schematic representation of the ATP-Insulin secretion model. Glucose is metabolized to ATP through glycolysis and oxidative phosphorylation. The increase in ATP triggers an influx of $Ca^{2+}$, which induces insulin release from the rapid release pool (RRP). Replenishment of the RRP is regulated by the intracellular glucose concentration. Created in BioRender. Henning, P. (2025) `https://BioRender.com/ifjt5zy`

#### 1.2.1 Species

| Species | Description |
|---|---|
| Cell(Gluc: float[mol/L]) | beta-cell, Gluc: glucose level |
| Mito | Mitochondrion |
| pyruvat_1000000 | $10^6$ pyruvate molecules |
| ATP_1000000 | $10^6$ ATP molecules |
| RRP_100 | 100 insulin molecules in the rapid releasable pool (RRP) |
| Ins_100 | 100 insulin molecules |
| Ca_100 | 100 $Ca^{2+}$ atoms |

**Table 4.** Species used in the ATP-Insulin secretion model.

#### 1.2.2 Rules

**Glycolysis**

For glycolysis, it is assumed that there is a constant supply of glucose, that metabolic activity does not lower the glucose level, and that glucose shuttling into the cell is not a bottleneck of the metabolism. Consequently, glucose is modeled as an attribute of the cell, rather than a species, and the glucose shuttling is not implemented. The ten-step glycolytic pathway is simplified to a single reaction that produces two pyruvate molecules and two ATP molecules, with ATP acting as a feedback inhibitor. NADH is not modeled as a species; however, for subsequent reactions, it is assumed that one NADH is produced per pyruvate molecule.

```
Cell{ATP_1000000:atp} -> Cell{2 pyruvat_1000000 + 3 ATP_1000000}
 @@ Vm_glyc*Cell.Gluc/((Km_glyc+Cell.Gluc)*(1+#atp/Ki_glyc));
```
(11)

This reaction is inhibited by ATP, consistent with regulatory control of key glycolytic enzymes [22].

**Anaerobic fermentation**

Pyruvate can be converted to lactate via anaerobic fermentation, consuming one NADH per pyruvate and thereby balancing the NADH generated in glycolysis.

```
pyruvat_1000000:pyr -> @@ Vmax_LDH*#pyr*1e6/(#pyr*1e6+Km_LDH_pyr);
```
(12)

**TCA cycle + oxidative phosphorylation**

Alternatively, pyruvate is oxidized in the tricarboxylic acid cycle and oxidative phosphorylation (oxPhos) to produce ATP within mitochondria.

```
Mito + pyruvat_1000000:pyr -> Mito + 15 ATP_1000000
 @@ (1/75)*Vmax_PDH*#pyr*1e6/(#pyr*1e6+Km_PDH_pyr);
```
(13)

In this reaction, NADH generated by glycolysis contributes to ATP production. The simplified pathway yields two ATP per glucose through glycolysis and fermentation, or 32 ATP per glucose through glycolysis and oxidative phosphorylation.

**ATP consumption**

ATP is consumed in various cellular processes in an ATP-dependent manner.

```
ATP_1000000:atp -> @@ k_ATP_u*#atp;
```
(14)

**$Ca^{2+}$ dynamic**

An increase in ATP concentration induces an influx of $Ca^{2+}$ ions. This process is represented by two rules controlling calcium increase and decrease depending on the ATP level:

```
Cell{Ca_100:ca+ATP_1000000:atp} -> Cell{Ca_100+Ca_100+ATP_1000000}
 @@ if (Ca_min + Ca_diff/(1+exp(Ca_rise*(-1*(#atp)+Ca_ATP))))>
 #ca then k_ca_flux;
```
(15)

```
Cell{Ca_100:ca+ATP_1000000:atp} -> Cell{ATP_1000000}
 @@ if (Ca_min + Ca_diff/(1+exp(Ca_rise*(-1*(#atp)+Ca_ATP))))<
 #ca then k_ca_flux;
```
(16)

This set of rules is a simplified version of a more complex process. Rising ATP concentration triggers the closure of ATP-sensitive potassium ($K^+$) channels, leading to a depolarization of the cell membrane. Due to the reduced cell membrane potential, voltage-gated calcium ($Ca^{2+}$) channels open, allowing calcium to enter the cell. However, for this model and the timescale of minutes/hours, we used simple rules that couples the ATP level to a $Ca^{2+}$ level.

**Insulin secretion**

The insulin secretion dynamics follow [23], where $Ca^{2+}$ triggers the release of insulin from the rapid release pool (RRP).

```
Cell -> Cell{RRP_100} @@ if ##(Ca_100 in Cell)<Ca_threshold then
 kr_gluc*Cell.Gluc+kr_offset else
 kr_gluc*Cell.Gluc+kr_offset+kr_ca*(##(Ca_100 in Cell)-Ca_threshold);
```
(17)

The glucose level in the cell controls RRP replenishment.

```
RRP_100:rrp -> Ins_100 @@ if ##(Ca_100 in Cell)<Ca_threshold then
 kf_ca1*(##(Ca_100 in Cell))*#rrp else
 kf_ca2*(##(Ca_100 in Cell))*#rrp;
```
(18)

#### 1.2.3 Initial conditions

| Species | Number | Ref. |
|---|---|---|
| Cell(Gluc = 2.5[millimol/L] | 1 | - |
| Mito | 75 | [12] |
| pyruvat_1000000 | 0 | - |
| ATP_1000000 | 4000 | [24] |
| RRP_100 | 0 | - |
| Ins_100 | 0 | - |
| Ca_100 | 500 | [17] |

**Table 5.** Initialization of the ATP-Insulin secretion submodel.

#### 1.2.4 Parameters

| Parameter | Description | Value | Ref. |
|---|---|---|---|
| vol_cell | Volume of a Min6 cell | 1025 $\mu^3 m^3$ | [19] |
| NA | Avogadro constant | 6.022 1/mol | - |
| Vm_glyc | Rate coefficient of glycolysis | 3408.964 1/min | fit to [24,25] |
| Km_glyc | Michaelis constant of glycolysis | 4.626 mM | [24] |
| Ki_glyc | Inhibition constant of glycolysis | 42806.181 | fit to [24,25] |
| Vmax_LDH | Rate coefficient of LDH | 5450 [1/min] | [26] |
| Km_LDH_pyr | Michaelis constant of LDH | 47.5 [$\mu$M]∗vol_cell ∗ NA | [27] |
| Vmax_PDH | Rate coefficient of oxPhos | 363.33 [1/min] | [25,26] |
| Km_PDH_pyr | Michaelis constant of oxPhos | 47.5 [$\mu$M]∗vol_cell ∗ NA | [27] |
| k_ATP_u | Rate coefficient of ATP use | 0.748 [1/min] | fit to [24,25] |
| Ca_min | Rate parameter of $Ca^{2+}$ flux | 486.4 | [17] |
| Ca_diff | Rate parameter of $Ca^{2+}$ flux | 742.5 | [17] |
| Ca_rise | Rate parameter of $Ca^{2+}$ flux | $0.759*10^{-3}$ | [17] |
| Ca_ATP | Rate parameter of $Ca^{2+}$ flux | 8719 | [17] |
| k_ca_flux | Rate parameter of $Ca^{2+}$ flux | 10 [1/s] | - |
| Ca_threshold | $Ca^{2+}$ act. threshold of GSIS | 852.936 | fit to [28] |
| kr_gluc | Rate parameter of GSIS | 261.542 [1/Ms] | fit to [28] |
| kr_offset | Rate parameter of GSIS | 4.796 [Hz] | fit to [28] |
| kr_ca | Rate parameter of GSIS | 0.103 [1/s] | fit to [28] |
| kf_ca1 | Rate parameter of GSIS | $0.125*10^{-2}$ [1/s] | fit to [28] |
| kf_ca2 | Rate parameter of GSIS | $0.326*10^{-2}$ [1/s] | fit to [28] |

**Table 6.** Parameters used in the submodel.

#### 1.2.5 Parameter Estimation

The submodel was used to estimate the ATP and insulin secretion dynamic parameters in three steps, as shown in the provenance graph in Fig.3 of the main paper. First, only the initial four rules describing ATP generation were used to fit the parameters Vm_glyc, Ki_glyc, and k_ATP_u to the glucose-dependent ATP production observed in Min6 cells. As fitting targets, both the intracellular ATP content at different glucose concentrations [24] and ATP production rates derived from the oxygen consumption rate (OCR) and extracellular acidification rate (ECAR) were used. The latter were calculated from the data reported by [25] using the equations provided in [29]. Parameter optimization was performed using pyABC, minimizing the relative quadratic error.

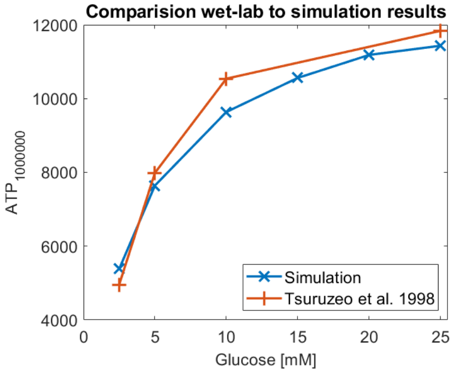


**Fig 5.** Comparison of ATP levels reported by [24] and the simulation results.

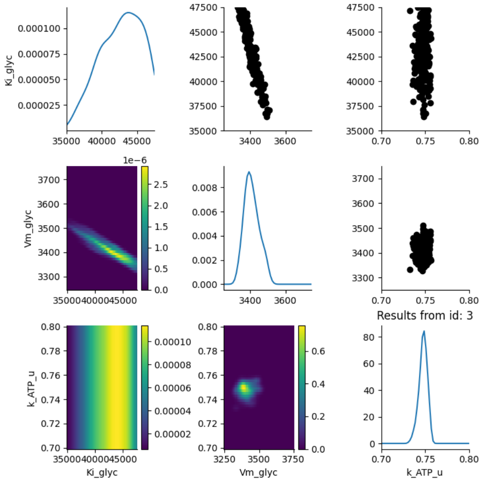


**Fig 6.** Posterior distributions of the fitted parameters obtained using pyABC.

The simulated ATP data show good agreement with the experimental measurements across different glucose levels (Fig. 5). Fig. 6 shows the posterior distributions of the three fitted parameters, all of which show convergence. After fitting the ATP dynamics, the calcium dynamics were added. The parameters `Ca_min`, `Ca_diff`, `Ca_rise`, and `Ca_ATP` from the rules 15 and 16 were estimated based on the calcium levels at different glucose levels reported in [17]. Finally, the parameters of the insulin secretion dynamics were estimated. The parameters in question were `Ca_threshold`, `kr_gluc`, `kr_offset`, `kr_ca`, `kf_ca1`, and `kf_ca2`. Parameter estimation was again performed using pyABC (with a relative quadratic error) based on the glucose-dependent insulin secretion data for Min6 cells [28].

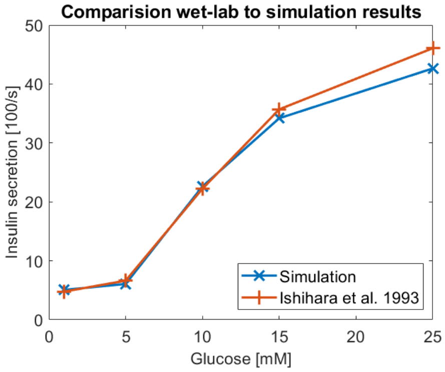


**Fig 7.** Comparison of the insulin secretion reported by [28] and the simulation model.

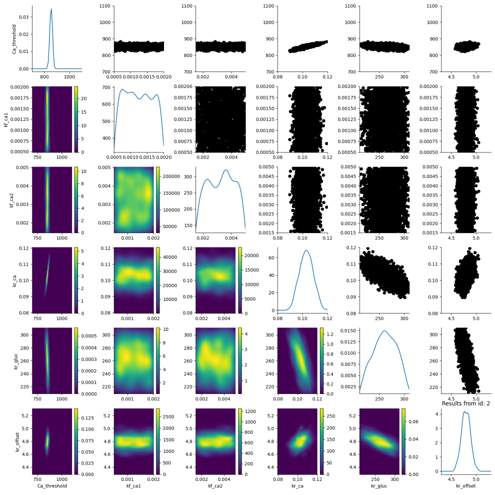

**Fig 8.** Posterior distributions of the fitted insulin secretion parameters obtained using pyABC.

The simulated insulin secretion dynamics show good agreement with experimental data (Fig. 7). Fig. 8 illustrates the posterior distributions of the six fitted parameters. It is notable that the distribution of the parameters `kf_ca1` and `kf_ca2` does not converge. These parameters are linked to the triggering pathway of insulin secretion but are insensitive to long-term behavior. Since the insulin secretion data from [28] (see fig. 7) only contains the accumulated insulin secretion over 2 hours and no time-resolved insulin secretion, these parameters have little effect on the simulation output and can not be determined by pyABC. To estimate these parameters, time-resolved data of the initial insulin secretion dynamics would be needed. Addressing this short-term dynamic, however, lies beyond the scope of the present study.

### 1.3 cAMP model

The cAMP submodel focuses on the dynamics of cyclic AMP (cAMP) generation and degradation shown in Fig. 9. This submodel uses parts of the previous model, as discussed below.

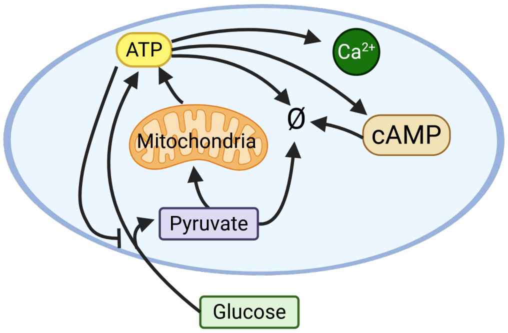


**Fig 9.** Schematic representation of the cAMP model. Glucose is metabolized to ATP, which is turned into cAMP by adenylyl cyclase (AC). cAMP decays over time. Created in BioRender. Henning, P. (2025) `https://BioRender.com/xooeky5`

#### 1.3.1 Species

| Species | Description |
|---|---|
| Cell(Gluc: float[mol/L]) | beta-cell, Gluc: glucose level |
| Mito | Mitochondrion |
| pyruvat_1000000 | $10^6$ pyruvate molecules |
| ATP_1000000 | $10^6$ ATP molecules |
| Ca_100 | 100 $Ca^{2+}$ atomes |
| cAMP_100 | 100 cAMP molecules |

**Table 7.** Species used in the cAMP model.

#### 1.3.2 Rules

The cAMP-submodel includes the first six rules (rules 11, 12, 13, 14, 15, and 16) of the previous model. Additionally, it includes a rule to model the generation of cAMP through adenylyl cyclase (AC) [30] and its subsequent degradation.

**cAMP generation**

As in other models [31], the rate of cAMP generation depends only on the intracellular calcium level [32,33], not on the ATP level. This assumption is supported by the fact that the $K_{m,ATP}$ value of AC (0.11 - 0.34 mM) [34,35] is much lower than the physiological ATP concentration in pancreatic beta cells (approximately 6–18 mM) [24]. The rate expression is derived from experimental data on AC activity at varying calcium levels [36].

```
Cell -> Cell{cAMP_100}
 @@ vmax_AC*((##(Ca_100 in Cell))^2)/(Km_Ca^2+((##(Ca_100 in Cell))^2));
```
(19)

**cAMP decay**
The degradation of cAMP is modeled as a first-order decay process with a constant rate [37,38].

```
Cell{cAMP_100} -> Cell @ k_cAMP_d;
```
(20)

#### 1.3.3 Initial conditions

| Species | Number | Ref. |
|---|---|---|
| Cell(Gluc = 2[millimol/L]) | 1 | - |
| Mito() | 75 | [12] |
| pyruvat_1000000 | 0 | - |
| ATP_1000000 | 4000 | [24] |
| Ca_100 | 500 | [17] |
| cAMP_100 | 6200 | [16] |

**Table 8.** Initialization of the cAMP submodel.

#### 1.3.4 Parameters

| Parameter | Description | Value | Ref. |
|---|---|---|---|
| vol_cell | Volume of a Min6 cell | 1025 $\mu^3 m^3$ | [19] |
| NA | Avogadro constant | 6.022 1/mol | - |
| Vm_glyc | Rate coefficient of glycolysis | 3408.964 1/min | see tab. 6 |
| Km_glyc | Michaelis constant of glycolysis | 4.626 mM | [24] |
| Ki_glyc | Inhibition constant of glycolysis | 42806.181 | see tab. 6 |
| Vmax_LDH | Rate coefficient of LDH | 5450 [1/min] | [26] |
| Km_LDH_pyr | Michaelis constant of LDH | 47.5 [$\mu$M]$*$vol_cell $*$ NA | [27] |
| Vmax_PDH | Rate coefficient of oxPhos | 363.33 [1/min] | [25,26] |
| Km_PDH_pyr | Michaelis constant of oxPhos | 47.5 [$\mu$M]$*$vol_cell $*$ NA | [27] |
| k_ATP_u | Rate coefficient of ATP use | 0.748 [1/min] | see tab. 6 |
| Ca_min | Rate parameter of $Ca^{2+}$ flux | 486.4 | [17] |
| Ca_diff | Rate parameter of $Ca^{2+}$ flux | 742.5 | [17] |
| Ca_rise | Rate parameter of $Ca^{2+}$ flux | $0.759*10^{-3}$ | [17] |
| Ca_ATP | Rate parameter of $Ca^{2+}$ flux | 8719 | [17] |
| k_ca_flux | Rate parameter of $Ca^{2+}$ flux | 10 [1/s] | - |
| vmax_AC | Rate coefficient of AC | 8000 [1/min] | - |
| k_cAMP_d | Decay of cAMP | 0.2602 [1/min] | fit to [16] |
| Km_Ca | Activation of AC | 1045 | fit to [16] |

**Table 9.** Parameters used in the submodel.

The parameter `vmax_AC` was set arbitrarily since the final level of cAMP depends on the ratio between the creation and decay of cAMP. Therefore, either `vmax_AC` or `k_cAMP_d` can be fixed to a sufficiently large value for cAMP to reach its steady state fast, and the other value can be used as a fit parameter to determine the steady state.

#### 1.3.5 Parameter Estimation

The cAMP submodel was used to estimate the parameters `k_cAMP_d` and `Km_Ca`, which describe cAMP decay and calcium-dependent activation, respectively. Experimental data from [16] were used as reference. Because quantitative cAMP measurements at multiple glucose levels in beta cells are scarce, the dataset includes cAMP concentrations at only two glucose concentrations. Parameter estimation was performed using pyABC, minimizing the relative quadratic error.

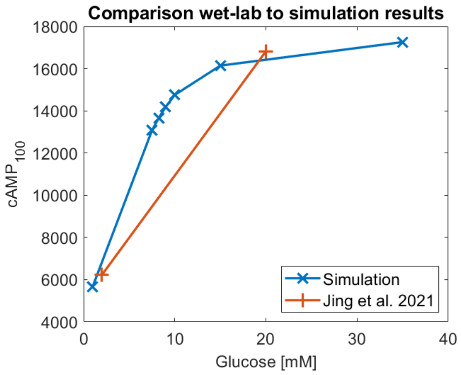


**Fig 10.** Comparison of the cAMP levels reported by [16] and simulation results from the cAMP submodel.

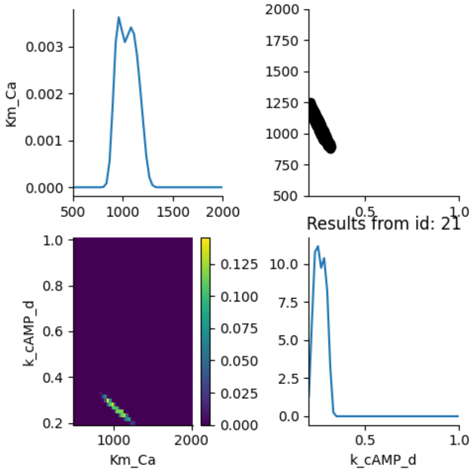


**Fig 11.** Posterior distributions of the fitted parameters obtained using pyABC.

Fig. 10 shows that the simulated cAMP levels are in good agreement with the experimental data. The narrow posterior peaks in Fig. 11 indicate that both parameters are well constrained and that the distributions have converged.

### 2 Beta-Cell Model

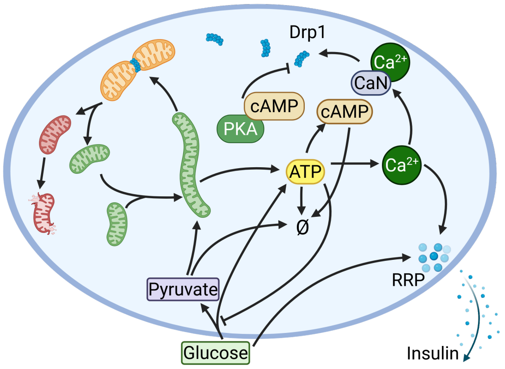


**Fig 12.** Schematic representation of the complete beta-cell model, integrating the ATP–insulin secretion, cAMP, and mitochondrial fission–fusion submodels. Created in BioRender. Henning, P. (2025) `https://BioRender.com/s00y554`

For the complete beta cell model (see Fig. 12), all three submodels were combined, and some extensions and modifications were added. Specifically, the anchor protein used in the fission-fusion submodel was replaced by the two distinct anchor proteins, MFF and Fis1. Additionally, mitochondrial compartments were extended to include the attributes `health` and `damage`, while the species `Mito_vol` used in the original fission–fusion model was converted into an attribute of each mitochondrion. For each 0.01 $\mu m^3$ of mitochondrial volume, one unit of `health` or `damage` is assigned to a mitochondrion. So in a mitochondrion with a size of 1 $\mu m^3$, a typicall size in Min6 cells [12], the health and damage attribute would add up to 100. To ensure that this ratio remains constant, it was necessary to model the volume as an attribute as well.

#### 2.1 Species

| Species | Description |
|---|---|
| Cell(Gluc: float[mol/L], mito_vol:int*) | beta-cell, Gluc: glucose level, mito_vol: volume of all mitochondria |
| Mito(vol:int*, health:int*, damage:int*, rand1:float*, rand2:float*) | Mitochondrion, vol: volume, health: health, damage: damage rand1/2: used for fission |
| Drp1_10(p:bool) | 10 Drp1 proteins, p: phosphorization state |
| Fis1 | Fis1 molecule |
| MFF | MFF molecule |
| PKA | cAMP-dependent protein kinase |
| cAMP | cyclic AMP |
| CaN | Calcineurin |
| Ca_100 | 100 $Ca^{2+}$ |
| pyruvat_1000000 | $10^6$ pyruvate molecules |
| ATP_1000000 | $10^6$ ATP molecules |
| RRP_100 | 100 insulin in rapid releasable pool (RRP) |
| Ins_100 | 100 insulin molecules |

**Table 10.** Species used in the beta-cell model.

### 2.2 Rules

**Drp1 binding**

Besides the changes from a general anchor protein to MFF and Fis1, the rules for Drp1 binding remain the same as rule 1.

```
Mito{Fis1} + Drp1_10(p==false) -> Mito{Fis1{Drp1_10}} @ kon/vol_cell;
Mito{MFF} + Drp1_10(p==false) -> Mito{MFF{Drp1_10}} @ kon/vol_cell;
```
(21)

**Drp1 unbinding**

The unbinding rules remain the same as in rule 2, besides the change of the anchor proteins.

```
Mito{Fis1:f{Drp1_10}} -> Mito{Fis1} + Drp1_10 @@ 1/10*koff*#f;
Mito{MFF:m{Drp1_10}} -> Mito{MFF} + Drp1_10 @@ 1/10*koff*#m;
```
(22)

**Drp1 (de)phosphorylation**

The rules for the phosphorization of Drp1 by PKA (rule 3) and the dephosphorization by CaN (rule 4) remain unchanged.

**Mitochondrial fission**

The fission rules were extended by the redistribution of the health and damage attributes. For the midzone fission, health and fission are divided proportionally to the size of the new sister mitochondria.

```
Mito:m{MFF{Drp1_10:drp + ?solA}} ->
 Mito(rand1 = RandomGauss(0.5, 0.075)){MFF} + Drp1_10+?solA
 @@ if (#drp>N_fission) then k_fission*(#(Mito_vol in m))^3;
```
(23)

```
Mito(rand1<1000):m{?sol1[Mito.rand1] + ?sol2[1-Mito.rand1]} ->
 Mito(vol=ceil(m.vol*m.rand1), health=ceil(m.health*m.rand1),
 damage=ceil(m.damage*m.rand1), rand1=1001, rand2=1001){?sol1} +
 Mito(vol=floor(m.vol*(1-m.rand1)), health = floor(m.health*(1-m.rand1)),
 damage = floor(m.damage*(1-m.rand1)), rand1=1001, rand2=1001){?sol2}
 @@ immediately;
```
(24)

In the peripheral fission event, the smaller mitochondria receive a disproportionately high amount of damage, controlled by the parameter `asym_fission`.

```
Mito:m{Fis1{Drp1_10:drp + ?solA}} ->
 Mito(rand2 = RandomGauss(0.2, 0.04)){Fis1}+Drp1_10+?solA
 @@ if (#drp>N_fission) then k_fission*(m.vol)^3;
```
(25)

```
Mito(rand2<1000, rand2>asym_fission, health+damage>=2):m
{?sol1[Mito.rand2] + ?sol2[1-Mito.rand2]} ->
 Mito(vol=ceil(m.vol*m.rand2), health=ceil(m.health*m.rand2-
 m.damage*asym_fission), damage=ceil(m.damage*(m.rand2+asym_fission)),
 rand1=1001, rand2=1001){?sol1} +
 Mito(vol=floor(m.vol*(1-m.rand2)), health=floor(m.health*(1-m.rand2)+
 m.damage*asym_fission), damage=floor(m.damage*((1-m.rand2)-asym_fission)),
 rand1=1001, rand2=1001){?sol2} @@ immediately;
```
(26)

In addition, a rule was added to the peripheral fission that prevents the creation of mitochondria with negative health.

```
Mito(rand2<=asym_fission) -> Mito(rand2=asym_fission + 0.0001)
@@ immediately;
```
(27)

**Mitochondrial Fusion**
In a fusion event, the health, damage, volume, and content of the two fusion mitochondria are combined in newly formed mitochondrion. This reaction can only occur, if the damage of the mitochondria is below the threshold `Thr_damage` [39].

```
Mito:m1{?solm1} + Mito:m2{?solm2} ->
 Mito(vol=m1.vol+m2.vol, health=m1.health+m2.health,
 damage=m1.damage+m2.damage, rand1=1001, rand2=1001){?solm1 + ?solm2}
 @@ if ((m1.damage/(m1.health+m1.damage))<Thr_damage &&
 (m2.damage/(m2.health+m2.damage))<Thr_damage) then k_fusion;
```
(28)

**ATP dynamic**
For the ATP generate the rules 11, 12, 14 remain the same. The reaction rate of oxidative phosphorylation is modified by a factor that considers that mitochondria with low health are less active [40–45]. See also eq. 5 of the main text.

```
Mito:m + pyruvat_1000000:pyr -> Mito + 15 ATP_1000000 @@
 f_ATP*(0.1+0.9/(1+exp(damage_m*(damage_lim-m.health/(m.health+
 m.damage)))))*(m.vol/N0)*Vmax_PDH*#pyr*1e6/(#pyr*1e6+Km_PDH_pyr);
```
(29)

**$Ca^{2+}$, cAMP, and insulin dynamic**
The rules for the $Ca^{2+}$ dynamic (rules 15 and 16), the cAMP dynamics (rules 19 and 20), and the insulin secretion (rules 17 and 18) are the same as in the submodel from section 1.2.

**Mitochondrial damage**
The primary addition in the comprehensive beta cell model is the representation of damage dynamics. It consists of the damage dealt to mitochondria during their metabolic activity [46].

```
Mito(health>0):m + pyruvat_1000000:pyr ->
 Mito(damage = m.damage+1, health = m.health-1) + pyruvat_1000000 @@
 f_ATP*(1+0.5*m.damage/(m.health+m.damage))*(0.1+0.9/(1+exp(damage_m*
 (damage_lim-m.health/(m.health+m.damage)))))*(m.vol/N0)*0.007*
 Vmax_PDH*#pyr*1e6/(#pyr*1e6+Km_PDH_pyr);
```
(30)

**Mitochondrial repair**
Parts of this damage can be repaired by the turnover of proteins [47–49] or mtDNA [40, 47].

```
Mito(damage>0):m -> Mito(damage = m.damage-1, health = m.health+1)
 @@ k_repair*m.damage;
```
(31)

**Mitophagy**

The repair rate is not sufficient to compensate for all the damage caused by the oxPhos. Therefore, damage accumulates over time in the mitochondrial network. This damage is removed by the removal of (sufficiently small) mitochondria with a large amount of damage [49].

```
Cell:c{Mito:m{?solm}} -> Cell(mito_vol=c.mito_vol-m.vol){?solm}
 @@ if (m.vol < 50 && m.damage/(m.damage+m.health) > Thr_damage)    (32)
 then 10 [Hz];
```

If a mitochondrion is removed, the MFF and Fis1 anchor sites bind to a new mitochondrion, holding the amount of Fis1 and MFF anchor sites constant.

```
Cell{Mito:m + Fis1} -> Cell{Mito{Fis1}} @@ 1000 [Hz]*(m.vol);
Cell{Mito:m + MFF} -> Cell{Mito{MFF}} @@ 1000 [Hz]*(m.vol);          (33)
```

**Biogenesis**
To compensate for the loss of mitochondrial mass through mitophagy, a biogenesis rule is introduced that enables mitochondria to grow.

```
Cell:c{Mito:m} -> Cell(mito_vol=c.mito_vol+1){Mito(vol = m.vol+1,}
 health = m.health+1) @@ k_biog*N0/(1+exp((c.mito_vol-(N0-100))/gamma));
                                                                      (34)
```

### 2.3 Initial conditions

| Species | Number | Ref. |
|---|---|---|
| Cell(Gluc = 1[milli mol/L], mito_vol = 6800) | 1 | - |
| Mito(vol = 100, health = 70, damage = 30, rand1 = 1001, rand2 = 1001) | 68 | [12] |
| Drp1_10(p = true) | 10000 | [13] |
| Drp1_10(p = false) | 10000 | [13] |
| Fis1 | 306 | [14, 15] |
| MFF | 306 | [14, 15] |
| PKA | 300000 | [13] |
| cAMP | 6200 | [16] |
| CaN | 350000 | [13] |
| Ca_100 | 500 | [17] |
| pyruvat_1000000 | 0 | - |
| ATP_1000000 | 4000 | [24] |
| RRP_100 | 0 | - |
| Ins_100 | 0 | - |

**Table 11.** Initialization of the cAMP submodel.

#### 2.4 Parameters

| Parameter | Description | Value | Ref. |
|---|---|---|---|
| vol_cell | Volume of a Min6 cell | 1025 $\mu m^3$ | [19] |
| NA | Avogadro constant | 6.022 1/mol | - |
| kon | Binding rate or Drp1 | 0.025 $\mu m^3/s$ | fit to [3,10,12,18] |
| Kd | Equi. constant for Drp1 binding | 0.301 $\mu mol/s$ | fit to [3,10,12,18] |
| koff | Unbinding rate of Drp1 | $\texttt{kon} * \texttt{Kd}$ | - |
| k_fission | Rate coefficient for fission | $8.211 \cdot 10^{-8} Hz$ | fit to [3,10,12,18] |
| N_fission | Number of Drp1 needed for fission | 11 | [3] |
| rand1 | Position of midzone fission | N(0.5, 0.075) | [9] |
| rand2 | Position of peripheral fission | N(0.2, 0.004) | [9] |
| asym_fission | Fission asymmetry | 0-10% | - |
| k_fusion | Rate coefficient for fusion | $5.41 \cdot 10^{-4}$ 1/min | [10,18] |
| kcatCaN | Rate constant for CaN | 7.6 $1/s$ | [7] |
| KMCaN | Michaelis constant for CaN | 160 mM | [7] |
| K_act_CaN | Activation constant of CaN | 14 nM | [8] |
| kcatpka | Rate constant ok PKA | 4 1/s | [20] |
| KMpka | Michaelis constant for PKA | 70 mM | [20] |
| K_act_PKA | Activation constant of PKA | 0.9 $\mu M$ | [5] |
| Vm_glyc | Rate coefficient of glycolysis | 3408.964 1/min | fit to [24,25] |
| Km_glyc | Michaelis constant of glycolysis | 4.626 mM | [24] |
| Ki_glyc | Inhibition constant of glycolysis | 42806.181 | fit to [24,25] |
| Vmax_LDH | Rate coefficient of LDH | 5450 [1/min] | [26] |
| Km_LDH_pyr | Michaelis constant of LDH | 47.5 [$\mu$M]$\cdot \texttt{vol\_cell} \cdot \texttt{NA}$ | [27] |
| Vmax_PDH | Rate coefficient of oxPhos | 363.33 [1/min] | [25,26] |
| Km_PDH_pyr | Michaelis constant of oxPhos | 47.5 [$\mu$M]$\cdot \texttt{vol\_cell} \cdot \texttt{NA}$ | [27] |
| f_ATP | Scaling factor for oxPhos | 1.54 | fit to [18,28] |
| damage_m | Steepness of damage response | 28.7 | fit to [18,28] |
| damage_lim | Point of damage response | 53.2% | fit to [18,28] |
| k_ATP_u | Rate coefficient of ATP use | 0.748 [1/min] | fit to [24,25] |
| Ca_min | Rate parameter of $Ca^{2+}$ flux | 486.4 | [17] |
| Ca_diff | Rate parameter of $Ca^{2+}$ flux | 742.5 | [17] |
| Ca_rise | Rate parameter of $Ca^{2+}$ flux | $0.759 \cdot 10^{-3}$ | [17] |
| Ca_ATP | Rate parameter of $Ca^{2+}$ flux | 8719 | [17] |
| k_ca_flux | Rate parameter of $Ca^{2+}$ flux | 10 [1/s] | - |
| Ca_threshold | $Ca^{2+}$ act. threshold of GSIS | 852.936 | fit to [28] |
| kr_gluc | Rate parameter of GSIS | 261.542 [1/Ms] | fit to [28] |
| kr_offset | Rate parameter of GSIS | 4.796 [Hz] | fit to [28] |
| kr_ca | Rate parameter of GSIS | 0.103 [1/s] | fit to [28] |
| kf_ca1 | Rate parameter of GSIS | $0.125 \cdot 10^{-2}$ [1/s] | fit to [28] |
| kf_ca2 | Rate parameter of GSIS | $0.326 \cdot 10^{-2}$ [1/s] | fit to [28] |
| vmax_AC | Rate coefficient of AC | 8000 [1/min] | - |
| k_cAMP_d | decay of cAMP | 0.2602 [1/min] | fit to [16] |
| Km_Ca | Activation of AC | 1045 | fit to [16] |
| N0 | Normal mitochondiral mass | 6800 | [12] |
| gamma | Parameter of biogenesis | 15 | - |
| k_biog | Rate parameter of biogenesis | 0.05 [1/day] | [48–50] |
| k_repair | Rate or repair | 0.1 [1/day] | [48–50] |
| Thr_damage | Threshold for fission and mitophagy | 15-85% | - |

**Table 12.** Parameters used in the beta cell model.

### 2.5 Parameter Estimation

The model was used to estimate parameters for the damage regulation. Details are described in the section *Interplay of Mitochondrial Damage and Insulin Secretion* of the main text.

In addition, we tested whether the complete beta cell model still reproduces the data used for parameter estimation in the submodels. For the simulation, the parameters from Table 1 of the main text were used.

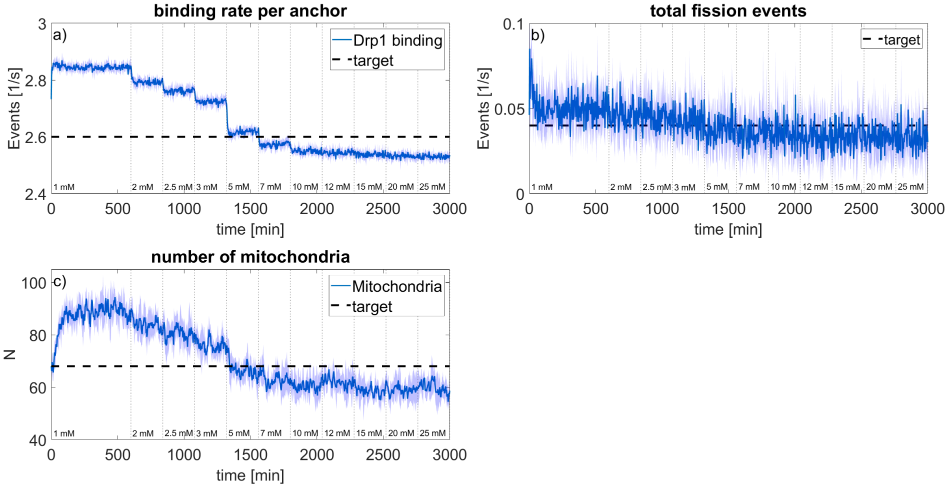


**Fig 13.** Comparison between literature data and simulation results for a) the Drp1 binding rate [3], b) the mitochondrial fission rate [10, 18], and c) the number of mitochondria in a Min6 cell [12]. The shaded area shows the standard derivation of N=5 simulation runs and the numbers at the bottom note the glucose level.

The output of the fission fusion dynamic is shown in Fig. 13. Initially, the model runs for 10 hours at 1 mM glucose to reach its steady state. After the steady state was achieved, the glucose level was increased every 4 hours. All observed parameters change in relation to the glucose level. For the Drp1 binding rate simulation results are within the range of the reported rate (2.6±1.7 1/s [3]). The number of mitochondria also lies within the range reported in the literature (50-100 mitochondria [12]). The fission rate varies slightly more than the reported values(0.039±0.006 fusion/mitochondrion/min [10] and 0.0457 fusion/mitochondrion/min, no standard deviation reported [18]); however, in the wet-lab experiments, the glucose concentration was not changed.

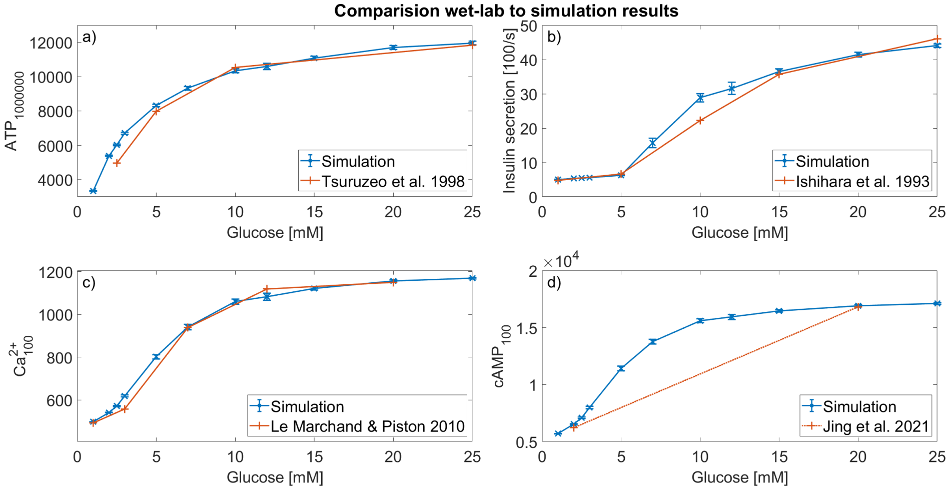


**Fig 14.** Comparison between the experimental and simulated data for a) ATP concentration, b) insulin secretion, c) calcium concentration, and d) cAMP concentration for different glucose levels. N=5 replications.

The simulated ATP, calcium, and cAMP concentrations, as well as the insulin secretion shown in Fig. 14 are still in good agreement in the complete beta cell model.
