## Appending and model files. for "Illuminating the Role of Asymmetric Mitochondrial Fission on Beta-Cell Health": S6_file.py

```
# -*- coding: utf-8 -*-
"""
Created on Thu May 16 09:52:16 2024

@author: phenning
"""

import numpy as np
import os
import pandas as pd
from timeit import default_timer as timer
import subprocess
import random
import string
import glob
import matplotlib.pyplot as plt
from pyswarms.single.global_best import GlobalBestPSO
from multiprocessing import Pool
from pyswarms.utils.plotters import (plot_cost_history, plot_contour, plot_surface)

# def simulation model
baseline = """
/* fission/fusion submodel */

/* ----- rate constants ----- */
const kon =  [micro^3 m^3/s];
const Kd =  [micro mol/L]*NA;
const k_fission = [Hz];

const koff = kon*Kd;
const N_fission = 11;
const k_fusion = 5.41e-4[1/min];

const vol_cell = 1025 [micro^3 m^3];

const kcatCaN = 7.6 [1/s];
const KMCaN = 160 [micro mol/L];
const K_act_CaN = 14e-9[mol/L]*vol_cell*NA;

const kcatpka= 4 [1/s];
const KMpka = 70 [micro mol/L];
const K_act_PKA = 0.9e-6[mol/L]*vol_cell*NA;
const NA = 6.022e23 [1/mol];

Cell();
Anchor();
Drp1_10(p: bool);
PKA();
CaN();
Ca_100();
cAMP_100();
Mito(rand1:float*, rand2:float*);
Mito_vol();

/* ----- Initial solution ----- */
Initial: Cell{68 Mito(rand1=-1, rand2=-1){9 Anchor{5 Drp1_10(p=false)} + 100 Mito_vol}  + 6940 Drp1_10(p=false) + 10000 Drp1_10(p=true) + 300000 PKA() + 350000 CaN() + 500 Ca_100() + 10000 cAMP_100()};

/* ----- Rules ----- */
/* Drp1_10 binding */
Mito{Anchor} + Drp1_10(p==false) -> Mito{Anchor{Drp1_10}} @ kon/vol_cell;

/* Drp1_10 unbinding */
Mito{Anchor:a{Drp1_10}} -> Mito{Anchor} + Drp1_10 @@ 1/10*koff*#a;

/* Midpoint fission */
Mito:m{Anchor{Drp1_10:drp + ?solA}} -> Mito(rand1 = RandomGauss(0.5, 0.075)){Anchor}+Drp1_10+?solA @@ if (#drp>N_fission) then k_fission*(#(Mito_vol in m))^3;
Mito(rand1>0):m{?sol1[Mito.rand1] + ?sol2[1-Mito.rand1]} -> Mito(rand1=-1, rand2=-1){?sol1} + Mito(rand1=-1, rand2=-1){?sol2} @@ immediately;

/* Peripheral Fission */
Mito:m{Anchor{Drp1_10:drp + ?solA}} -> Mito(rand2 = RandomGauss(0.2, 0.04)){Anchor}+Drp1_10+?solA @@ if (#drp>N_fission) then k_fission*(#(Mito_vol in m))^3;
Mito(rand2>0):m{?sol1[Mito.rand2] + ?sol2[1-Mito.rand2]} -> Mito(rand1=-1, rand2=-1){?sol1} + Mito(rand1=-1, rand2=-1){?sol2} @@ immediately;

/* fusion */
Mito:m1{?solm1} + Mito:m2{?solm2} -> Mito(rand1=-1, rand2=-1){?solm1 + ?solm2} @@ k_fusion;

/* dephosphorylation by CaN */
Cell{Drp1_10(p==true):d + CaN:c} -> Cell{Drp1_10(p=false) + CaN} @@ 1/10*kcatCaN*#c*#d*10/(KMCaN*vol_cell*NA+#d*10)*100*##(Ca_100 in Cell)/(K_act_CaN+100*##(Ca_100 in Cell));

/* phosphorylation by PKA */
Cell{Drp1_10(p==false):d + PKA:p} -> Cell{Drp1_10(p=true) + PKA} @@ 1/10*kcatpka*#p*#d*10/(KMpka*vol_cell*NA+#d*10)*100*##(cAMP_100 in Cell)/(K_act_PKA+100*##(cAMP_100 in Cell));

output ##Mito;
"""

# executes model in parallel
def run_par(rates):
    rates = rates.tolist()
    results = []
    pool = Pool()
    res = pool.map(model, rates)
    results.append(res)
    pool.close()
    pool.join()
    return np.array(results[0])

# exectes model one at a time
def run_seq(rates):
    n_particles = rates.shape[0]
    j = [model(rates[i,:]) for i in range(n_particles)]
    return np.array(j)

# added rates to the base model and execute it
def model(rates, t_end=3, Nt=300, N=1):
    silent = True
    
    # adds fit parameter to basemodel
    kon = float(rates[0])
    if kon == None:
        raise ValueError('No kon specified')
    model = baseline.replace("const kon = ","const kon = " + str(kon))
    
    Kd = float(rates[1])
    if Kd == None:
        raise ValueError('No Kd specified')        
    model = model.replace("const Kd = ","const Kd = " + str(Kd))

    k_fission = float(rates[2])
    if k_fission == None:
        raise ValueError('No k_fission specified')        
    model = model.replace("const k_fission = ","const k_fission = " + str(k_fission))


    # creates model file with random name
    random_string = "_"+''.join(random.choice(string.ascii_lowercase) for i in range(25))
    text_file = open("/home/phenning/pySwarm/Fit_DRP/model"+random_string+".mlr", "w")
    text_file.write(model)
    text_file.close()
    
    # runs ML-Rules3 model
    if silent:
        subprocess.check_output('cargo run --release --manifest-path /home/phenning/ml-rules-3/Cargo.toml -- /home/phenning/pySwarm/Fit_DRP/model'+random_string+'.mlr --simulate-until "'+str(t_end)+'[h]" --num-observation '+str(Nt)+' --dynamic-silent --output-prefix /home/phenning/pySwarm/Fit_DRP/output/output'+random_string+' --num-reps '+str(N)+' --output-rule-counts --output-csv',shell=True,stderr=subprocess.STDOUT)
    else:
        subprocess.check_output('cargo run --release --manifest-path /home/phenning/ml-rules-3/Cargo.toml -- /home/phenning/pySwarm/Fit_DRP/model'+random_string+'.mlr --simulate-until "'+str(t_end)+'[h]" --num-observation '+str(Nt)+' --dynamic-silent --output-prefix /home/phenning/pySwarm/Fit_DRP/output/output'+random_string+' --num-reps '+str(N)+' --output-rule-counts --output-csv',shell=True)
    
    # load output data
    if N != 1:
        df = pd.read_csv('/home/phenning/pySwarm/Fit_DRP/output/output'+random_string+'_0_rule_counts.csv', sep=',')
    else:
        df = pd.read_csv('/home/phenning/pySwarm/Fit_DRP/output/output'+random_string+'_rule_counts.csv', sep=',')
    time = np.unique(df['time[s]'])
    
    Drp_on = np.zeros((1, N))
    split = np.zeros((1, N))
    
    # calculate Drp1 binding events per anchor per second and split events per second
    for n in range(N):
        if N != 1:
            df = pd.read_csv('/home/phenning/pySwarm/Fit_DRP/output/output'+random_string+'_'+ str(n) +'_rule_counts.csv', sep=',')
        else:
            df = pd.read_csv('/home/phenning/pySwarm/Fit_DRP/output/output'+random_string+'_rule_counts.csv', sep=',')

        tmp = df['Mito{Anchor} + Drp1_10(p==false) -> Mito{Anchor{Drp1_10}} @ kon/vol_cell']
        Drp_on[:,n] = (tmp[len(tmp)-1]-tmp[200])/(time[-1]-time[200])*10/612 # Drp1 binding per anchor per second

        tmp1 = df['Mito(rand1>0):m{?sol1[Mito.rand1] + ?sol2[1-Mito.rand1]} -> Mito(rand1=-1  rand2=-1){?sol1} + Mito(rand1=-1  rand2=-1){?sol2} @@ immediately']
        tmp2 = df['Mito(rand2>0):m{?sol1[Mito.rand2] + ?sol2[1-Mito.rand2]} -> Mito(rand1=-1  rand2=-1){?sol1} + Mito(rand1=-1  rand2=-1){?sol2} @@ immediately']
        split[:,n] = ((tmp1[len(tmp1)-1]-tmp1[200])/(time[-1]-time[200]) + (tmp2[len(tmp2)-1]-tmp2[200])/(time[-1]-time[200]))*60/68 # Split events per min per mito

    output = np.mean(Drp_on)
    output2 = np.mean(split)

    Mitos = np.zeros((1, N))
    for n in range(N):
        if N != 1:
            df = pd.read_csv('/home/phenning/pySwarm/Fit_DRP/output/output'+random_string+'_'+ str(n) +'.csv', sep=',')
        else:
            df = pd.read_csv('/home/phenning/pySwarm/Fit_DRP/output/output'+random_string+'.csv', sep=',')
        tmp = df['##Mito']
        Mitos[:,n] = np.mean(tmp[200:-1])

    output3 = np.mean(Mitos)
    # deletes output files
    for filename in glob.glob('/home/phenning/pySwarm/Fit_DRP/output/*'+random_string+'*'):
        os.remove(filename) 
    for filename in glob.glob('/home/phenning/pySwarm/Fit_DRP/*'+random_string+'*'):
        os.remove(filename)
        
    # Drp_on data from Michalska et al. 2018 in [1/s]
	# split data from Twig et al. 2008 and Cagalinec et al. 2013 in [event/mito/min]
	# Mitos data from Chaudhry et al. 2020.
    data  = {'Drp_on': 2.6, 'split': 0.04, 'Mitos': 68}
	
	# calcules (relative) mean square error
    error = (
        pow((output - data['Drp_on'])/data['Drp_on'],2) + pow((output2 - data['split'])/data['split'],2) + pow((output3-data['Mitos'])/data['Mitos'],2)
    )
    return error
    
    
if __name__ == '__main__':
    # [kon, Kd, k_fission]
    min_bound = np.array([0.02, 0.2, 1e-8])
    max_bound = np.array([0.04, 0.4, 1e-7])
    bounds = (min_bound, max_bound)

    options = {'c1': 0.5, 'c2': 0.3, 'w': 0.9}
    optimizer = GlobalBestPSO(n_particles=256, dimensions=3, options=options, bounds=bounds, oh_strategy={"w":'exp_decay', 'c1':'lin_variation'})
    cost, pos = optimizer.optimize(run_par, 40)
    
    parameter = ['kon', 'Kd', 'k_fission']
    num = '1'
    log = open("/home/phenning/pySwarm/Fit_DRP/log_"+str(num)+".txt", "w")
    for i in range(0, len(pos)):
        log.write(str(parameter[i]) + " = " + str(pos[i]) + ", min = " + str(min_bound[i]) + ", max = " + str(max_bound[i]) + "\n")
    log.write("final cost value: " + str(cost))
    log.close()
    
    plot_cost_history(cost_history=optimizer.cost_history)
    plt.savefig('cost_DRP_fit_'+str(num)+'.png')
```
