## Appending and model files. for "Illuminating the Role of Asymmetric Mitochondrial Fission on Beta-Cell Health": S7_file.py

```
# -*- coding: utf-8 -*-
"""
Created on Thu May 16 09:52:16 2024

@author: phenning
"""

import numpy as np
import os
import pandas as pd
from timeit import default_timer as timer
import subprocess
import pyabc
import random
import string
import glob
import matplotlib.pyplot as plt


# def simulation model
baseline = """
/* ATP submodel */

/* ----- rate constants ----- */
const vol_cell = 1025 [micro^3 m^3];
const NA = 6.022e23 [1/mol];

const Vm_glyc = 2725 [1/min];
const Km_glyc = 4.626[milli mol/L];
const Ki_glyc = 4000;

const Vmax_LDH = 5450 [1/min];
const Km_LDH_pyr = 47.5 [micro mol/L]*vol_cell*NA;

const Vmax_PDH = 363.333 [1/min];
const Km_PDH_pyr = 47.5 [micro mol/L]*vol_cell*NA;

const k_ATP_u = 4 [1/min];

/* ----- Species ----- */
Cell(Gluc: float[mol/L]);
Mito();
pyruvat_1000000();
ATP_1000000();


/* ----- Initial solution ----- */
Initial: 1 Cell(Gluc = 2.5[milli mol/L]){75 Mito + 4000 ATP_1000000};

/* ----- Rules ----- */
/* glycolysis of 1000000 glucose molecule*/
Cell{ATP_1000000:atp} -> Cell{2 pyruvat_1000000 + 3 ATP_1000000} @@ Vm_glyc*Cell.Gluc/((Km_glyc+Cell.Gluc)*(1+#atp/Ki_glyc));

/* anerobmic fermentaion */
pyruvat_1000000:pyr -> @@ Vmax_LDH*#pyr*1e6/(#pyr*1e6+Km_LDH_pyr);

/* TCA + OxPhos*/
Mito + pyruvat_1000000:pyr -> Mito + 15 ATP_1000000 @@ (1/75)*Vmax_PDH*#pyr*1e6/(#pyr*1e6+Km_PDH_pyr);

/* ATP usage */
ATP_1000000:atp ->  @@ k_ATP_u*#atp;


/* ----- Glucose rise ------ */
Cell -> Cell(Gluc = 5 [milli mol/L]) @@ if time() > 4[h] && time() < 4.1[h] && Cell.Gluc<5 [milli mol/L] then immediately;
Cell -> Cell(Gluc = 10 [milli mol/L]) @@ if time() > 8[h] && time() < 8.1[h] && Cell.Gluc<10 [milli mol/L] then immediately;
Cell -> Cell(Gluc = 25 [milli mol/L]) @@ if time() > 12[h] && time() < 12.1[h] && Cell.Gluc<25 [milli mol/L] then immediately;
Cell -> Cell(Gluc = 20 [milli mol/L]) @@ if time() > 16[h] && time() < 16.1[h] && Cell.Gluc>20 [milli mol/L] then immediately;

output ##(ATP_1000000);
"""

# added rates to the base model and execute it
def model(rates, t_end=20, Nt=100, N=1):
    silent = True

    # adds fit parameter to basemodel
    k_ATP_u = rates['k_ATP_u']
    if k_ATP_u == None:
        raise ValueError('No k_ATP_u specified')
    model = baseline.replace("const k_ATP_u = 4","const k_ATP_u = " + str(k_ATP_u))
	
    Ki_glyc = rates['Ki_glyc']
    if Ki_glyc == None:
        raise ValueError('No Ki_glyc specified')        
    model = model.replace("const Ki_glyc = 4000","const Ki_glyc = " + str(Ki_glyc))
	
    Vm_glyc = rates['Vm_glyc']
    if Vm_glyc == None:
        raise ValueError('No Vm_glyc specified')        
    model = model.replace("const Vm_glyc = 2725","const Vm_glyc = " + str(Vm_glyc))
    
    # creates model file with random name    
    random_string = "_"+''.join(random.choice(string.ascii_lowercase) for i in range(25))
    text_file = open("/home/phenning/Documents/pyABC_Mito/Fit_ATP/model"+random_string+".mlr", "w")
    text_file.write(model)
    text_file.close()

    # runs ML-Rules3 model
    if silent:
            subprocess.check_output('cargo run --release --manifest-path /home/phenning/ml-rules-3/Cargo.toml -- /home/phenning/Documents/pyABC_Mito/Fit_ATP/model'+random_string+'.mlr --simulate-until "'+str(t_end)+'[h]" --num-observation '+str(Nt)+' --dynamic-silent --output-prefix /home/phenning/Documents/pyABC_Mito/Fit_ATP/output/output'+random_string+' --num-reps '+str(N)+' --output-csv --output-rule-counts', shell=True,stderr=subprocess.STDOUT)
    else:
        subprocess.check_output('cargo run --release --manifest-path /home/phenning/ml-rules-3/Cargo.toml -- /home/phenning/Documents/pyABC_Mito/Fit_ATP/model'+random_string+'.mlr --simulate-until "'+str(t_end)+'[h]" --num-observation '+str(Nt)+' --dynamic-silent --output-prefix /home/phenning/Documents/output/output'+random_string+' --num-reps '+str(N)+' --output-csv --output-rule-counts', shell=True)

    # load output data
    if N != 1:
        df = pd.read_csv('/home/phenning/Documents/pyABC_Mito/Fit_ATP/output/output'+random_string+'_0.csv', sep=',')
    else:
        df = pd.read_csv('/home/phenning/Documents/pyABC_Mito/Fit_ATP/output/output'+random_string+'.csv', sep=',')
    time = np.unique(df['time[s]'])
    
    
    Gluc = [2.5, 5, 10, 25]
    ATP = np.zeros((len(Gluc), N))
    
    # calculate mean ATP level
    for n in range(N):
        if N != 1:
            df = pd.read_csv('/home/phenning/Documents/pyABC_Mito/Fit_ATP/output/output'+random_string+'_'+ str(n) +'.csv', sep=',')
        else:
            df = pd.read_csv('/home/phenning/Documents/pyABC_Mito/Fit_ATP/output/output'+random_string+'.csv', sep=',')
        tmp = df['##(ATP_1000000)']
        for i in range(len(Gluc)):
            ATP[i,n] = (np.mean(tmp[np.logical_and(time>3600*(4*i+1), time<3600*(4*i+3))]))
    output = np.mean(ATP, 1)
    
    # calculate mean ATP usage
    ATP_use = np.zeros((N, 2))
    for n in range(N):
        if N != 1:
            df = pd.read_csv('/home/phenning/Documents/pyABC_Mito/Fit_ATP/output/output'+random_string+'_'+ str(n) +'_rule_counts.csv', sep=',')
        else:
            df = pd.read_csv('/home/phenning/Documents/pyABC_Mito/Fit_ATP/output/output'+random_string+'_rule_counts.csv', sep=',')
        tmp = df['ATP_1000000:atp -> @@ k_ATP_u*#atp']
        ATP_use[n, 0] = (tmp[(np.abs(time-3.5*3600)).argmin()]-tmp[(np.abs(time-0.5*3600)).argmin()])/(time[(np.abs(time-3.5*3600)).argmin()]-time[(np.abs(time-0.5*3600)).argmin()])
        ATP_use[n, 1] = (tmp[(np.abs(time-19.5*3600)).argmin()]-tmp[(np.abs(time-16.5*3600)).argmin()])/(time[(np.abs(time-19.5*3600)).argmin()]-time[(np.abs(time-16.5*3600)).argmin()])

    output2 = np.mean(ATP_use, 0)
    
    # deletes output files
    for filename in glob.glob('/home/phenning/Documents/pyABC_Mito/Fit_ATP/output/*'+random_string+'*'):
        os.remove(filename) 
    for filename in glob.glob('/home/phenning/Documents/pyABC_Mito/Fit_ATP/*'+random_string+'*'):
        os.remove(filename)
    return{"ATP":output, "ATP_use":output2}

# calcules (relative) mean square error
def distance(sim, data):
    error = (
        np.sum(pow((sim['ATP'] - data['ATP'])/data['ATP'],2)) + 10*np.sum(pow((sim['ATP_use'] - data['ATP_use'])/data['ATP_use'],2))
    )
    return error

def run_ABC():
    prior = pyabc.Distribution(
        k_ATP_u=pyabc.RV("uniform", 0.7, 0.1),
        Ki_glyc=pyabc.RV("uniform", 35000, 12500),
        Vm_glyc=pyabc.RV("uniform", 3250, 500))
        
    abc = pyabc.ABCSMC(
        models=model,
        parameter_priors=prior,
        distance_function=distance,
        population_size=pyabc.AdaptivePopulationSize(100, 0.14, 2500),
        sampler = pyabc.sampler.MulticoreEvalParallelSampler(n_procs=32),
    )
    
    # ATP data from Tsuruzeo et al. 1998 in [fmol / cell]
    # ATP usage data from Xiong et al. 2016 in [1e6 ATP / min]
    theo = {'ATP': np.array([8.23213871, 13.26282829, 17.50268618, 19.65686186])*1e-15*6.022e23/1000000, 'ATP_use': np.array([4125/60, 8416/60])}
    
    import tempfile
    db_path = "sqlite:///" + os.path.join(tempfile.gettempdir(), "/home/phenning/Documents/pyABC_Mito/Fit_ATP/pyABC_ATP_u.db")
    abc_id = abc.new(db_path, {"ATP": theo['ATP'], "ATP_use": theo['ATP_use']})
    
    history = abc.run(minimum_epsilon=0.01, max_nr_populations=6)
    return(abc_id, history, db_path)

def continue_ABC(model, prior, distance, db_path, abc_id):
    abc_continued = pyabc.ABCSMC(model,
        prior,
        distance,
        population_size=pyabc.AdaptivePopulationSize(100, 0.14, 2500),
        sampler = pyabc.sampler.MulticoreEvalParallelSampler(n_procs=32))
    abc_continued.load(db_path, abc_id)
    abc_continued.run(minimum_epsilon=0.001, max_nr_populations=4)
    
    return(abc_continued)
    
def plot_ABC(path, model, distance, id=1):
    prior = pyabc.Distribution(
        k_ATP_u=pyabc.RV("uniform", 0.7, 0.1),
        Ki_glyc=pyabc.RV("uniform", 35000, 12500),
        Vm_glyc=pyabc.RV("uniform", 3250, 500))
    abc_continued = pyabc.ABCSMC(model, prior, distance)
    abc_continued.load(path, id)
    history = abc_continued.history
    print('max_t = ', history.max_t)
    import matplotlib.pyplot as plt
    from pyabc.visualization import plot_kde_matrix
    fig = plt.figure(figsize=(10,8))
    dfw = history.get_distribution(m=0)
    plot_kde_matrix(
        *dfw,
        limits={"k_ATP_u":(0.7,0.8),
            "Ki_glyc":(35000,47500),
            "Vm_glyc":(3250,3750)})
    plt.title("Results from id: "+str(id))
    plt.savefig('k_ATP_id_'+str(id)+'.png')
    
def output_results(db_path):
    history = pyabc.History(db_path)
    df,w = history.get_distribution()
    names = list(df.columns.values)
    for i in names:
        print(i, " = ",pyabc.weighted_statistics.weighted_mean(np.array(df[i]),w))

if __name__ == '__main__':
	# ----- run pyABC ----- 
    abc_id, history, db_path = run_ABC()
    
    # ----- continu pyABC -----
    #import tempfile
    #db_path = "sqlite:///" + os.path.join(tempfile.gettempdir(), "/home/phenning/Documents/pyABC_Mito/Fit_ATP/pyABC_ATP_u.db")
    #prior = pyabc.Distribution(
    #    k_ATP_u=pyabc.RV("uniform", 0.7, 0.1),
    #    Ki_glyc=pyabc.RV("uniform", 35000, 12500),
    #    Vm_glyc=pyabc.RV("uniform", 3250, 500))
    #abc_continued = continue_ABC(model = model, prior = prior, distance = distance, db_path = db_path, abc_id = 3)
    
    # ----- plot results -----
    #import tempfile
    #path = "sqlite:///" + os.path.join(tempfile.gettempdir(), "/home/phenning/Documents/pyABC_Mito/Fit_ATP/pyABC_ATP_u.db")
    #plot_ABC(path = path, model = model, distance = distance, id = 3)
    
    # ----- output results -----
    #import tempfile
    #db_path = "sqlite:///" + os.path.join(tempfile.gettempdir(), "/home/phenning/Documents/pyABC_Mito/Fit_ATP/pyABC_ATP_u.db")
    #output_results(db_path)
    
```
