## Appending and model files. for "Illuminating the Role of Asymmetric Mitochondrial Fission on Beta-Cell Health": S8_file.py

```
# -*- coding: utf-8 -*-
"""
Created on Thu May 16 09:52:16 2024

@author: phenning
"""

import numpy as np
import os
import pandas as pd
from timeit import default_timer as timer
import subprocess
import pyabc
import random
import string
import glob
import matplotlib.pyplot as plt


# def simulation model
baseline = """
/* ATP-Ins submodel */

/* ----- rate constants ----- */
const vol_cell = 1025 [micro^3 m^3];
const NA = 6.022e23 [1/mol];

const Vm_glyc = 3408.964 [1/min];
const Km_glyc = 4.626 [milli mol/L];
const Ki_glyc = 42806.181;

const Vmax_LDH = 5450 [1/min];
const Km_LDH_pyr = 47.5 [micro mol/L]*vol_cell*NA;

const Vmax_PDH = 363.333 [1/min];
const Km_PDH_pyr = 47.5 [micro mol/L]*vol_cell*NA;

const vol_total = 75 [micro^3 m^3];

const k_ATP_u = 0.748 [1/min];

/* Ca2+ dynamic */
const Ca_min = 486.4;
const Ca_diff = 742.5;
const Ca_rise = 0.000759;
const Ca_ATP = 8719;
const k_ca_flux = 10 [1/s];

/* Insulin dynamic */
const Ca_threshold = ;
const kr_gluc =  [L/mol s];
const kr_offset =  [Hz];
const kr_ca =  [1/s];
const kf_ca1 =  [1/s];
const kf_ca2 =  [1/s];

/* ----- Species ----- */
Cell(Gluc: float[mol/L]);
Mito();
pyruvat_1000000();
ATP_1000000();
Ca_100();
RRP_100();
Ins_100();

/* ----- Initial solution ----- */
Initial: 1 Cell(Gluc = 1[milli mol/L]){75 Mito + 4000 ATP_1000000 + 500 Ca_100};

/* ----- Rules ----- */
/* glycolysis of 1000000 glucose molecule*/
Cell{ATP_1000000:atp} -> Cell{2 pyruvat_1000000 + 3 ATP_1000000} @@ Vm_glyc*Cell.Gluc/((Km_glyc+Cell.Gluc)*(1+#atp/Ki_glyc));

/* anerobmic fermentaion */
pyruvat_1000000:pyr -> @@ Vmax_LDH*#pyr*1e6/(#pyr*1e6+Km_LDH_pyr);

/* TCA + OxPhos*/
Mito + pyruvat_1000000:pyr -> Mito + 15 ATP_1000000 @@ (1/75)*Vmax_PDH*#pyr*1e6/(#pyr*1e6+Km_PDH_pyr);

/* ATP usage */
ATP_1000000:atp ->  @@ k_ATP_u*#atp;

/* Ca2+ influx */
Cell{Ca_100:ca+ATP_1000000:atp} -> Cell{Ca_100+Ca_100+ATP_1000000} @@ if (Ca_min + Ca_diff/(1+exp(Ca_rise*(-1*(#atp)+Ca_ATP))))>#ca then k_ca_flux;

/* Ca2+ outflux */
Cell{Ca_100:ca+ATP_1000000:atp} -> Cell{ATP_1000000} @@ if (Ca_min + Ca_diff/(1+exp(Ca_rise*(-1*(#atp)+Ca_ATP))))<#ca then k_ca_flux;

/* RRP + Insulin model taken from Grespan et al. 2017 */
/* filling of the readily releasable pool (RRL) */
Cell -> Cell{RRP_100} @@ if ##(Ca_100 in Cell)<Ca_threshold then kr_gluc*Cell.Gluc+kr_offset else kr_gluc*Cell.Gluc+kr_offset+kr_ca*(##(Ca_100 in Cell)-Ca_threshold);

/* Insulin secretion from RRP */
RRP_100:rrp -> Ins_100 @@ if ##(Ca_100 in Cell)<Ca_threshold then kf_ca1*(##(Ca_100 in Cell))*#rrp else kf_ca2*(##(Ca_100 in Cell))*#rrp;


/* ----- Glucose rise ------ */

Cell -> Cell(Gluc = 5 [milli mol/L]) @@ if time() > 2[h] && time() < 2.1[h] && Cell.Gluc<5 [milli mol/L] then immediately;
Cell -> Cell(Gluc = 10 [milli mol/L]) @@ if time() > 4[h] && time() < 4.1[h] && Cell.Gluc<10 [milli mol/L] then immediately;
Cell -> Cell(Gluc = 15 [milli mol/L]) @@ if time() > 6[h] && time() < 6.1[h] && Cell.Gluc<15 [milli mol/L] then immediately;
Cell -> Cell(Gluc = 25 [milli mol/L]) @@ if time() > 8[h] && time() < 8.1[h] && Cell.Gluc<25 [milli mol/L] then immediately;
Cell -> Cell(Gluc = 50 [milli mol/L]) @@ if time() > 10[h] && time() < 10.1[h] && Cell.Gluc<50 [milli mol/L] then immediately;


//output ##(ATP_1000000);
"""

# added rates to the base model and execute it
def model(rates, t_end=12, Nt=150, N=1):
    silent = True
    
    # adds fit parameter to basemodel
    Ca_threshold = rates['Ca_threshold']
    if Ca_threshold == None:
        raise ValueError('No Ca_threshold specified')
    model = baseline.replace("const Ca_threshold = ","const Ca_threshold = " + str(Ca_threshold))
    
    kr_gluc = rates['kr_gluc']
    if kr_gluc == None:
        raise ValueError('No kr_gluc specified')
    model = model.replace("const kr_gluc = ","const kr_gluc = " + str(kr_gluc))
    
    kr_offset = rates['kr_offset']
    if kr_offset == None:
        raise ValueError('No kr_offset specified')
    model = model.replace("const kr_offset = ","const kr_offset = " + str(kr_offset))
    
    kr_ca = rates['kr_ca']
    if kr_ca == None:
        raise ValueError('No kr_ca specified')
    model = model.replace("const kr_ca = ","const kr_ca = " + str(kr_ca))
    
    kf_ca1 = rates['kf_ca1']
    if kf_ca1 == None:
        raise ValueError('No kf_ca1 specified')
    model = model.replace("const kf_ca1 = ","const kf_ca1 = " + str(kf_ca1))
    
    kf_ca2 = rates['kf_ca2']
    if kf_ca2 == None:
        raise ValueError('No kf_ca2 specified')
    model = model.replace("const kf_ca2 = ","const kf_ca2 = " + str(kf_ca2))
    
    # creates model file with random name
    random_string = "_"+''.join(random.choice(string.ascii_lowercase) for i in range(25))
    text_file = open("/home/phenning/pyABC/Fit_Ins/model"+random_string+".mlr", "w")
    text_file.write(model)
    text_file.close()
    
    # runs ML-Rules3 model
    if silent:
            subprocess.check_output('cargo run --release --manifest-path /home/phenning/ml-rules-3/Cargo.toml -- /home/phenning/pyABC/Fit_Ins/model'+random_string+'.mlr --simulate-until "'+str(t_end)+'[h]" --num-observation '+str(Nt)+' --dynamic-silent --output-prefix /home/phenning/pyABC/Fit_Ins/output/output'+random_string+' --num-reps '+str(N)+' --output-csv --output-rule-counts', shell=True,stderr=subprocess.STDOUT)
    else:
        subprocess.check_output('cargo run --release --manifest-path /home/phenning/ml-rules-3/Cargo.toml -- /home/phenning/pyABC/Fit_Ins/model'+random_string+'.mlr --simulate-until "'+str(t_end)+'[h]" --num-observation '+str(Nt)+' --dynamic-silent --output-prefix /home/phenning/pyABC/Fit_Ins/output/output'+random_string+' --num-reps '+str(N)+' --output-csv --output-rule-counts', shell=True)

    # load output data
    if N != 1:
        df = pd.read_csv('/home/phenning/pyABC/Fit_Ins/output/output'+random_string+'_0.csv', sep=',')
    else:
        df = pd.read_csv('/home/phenning/pyABC/Fit_Ins/output/output'+random_string+'.csv', sep=',')
    time = np.unique(df['time[s]'])
    
    Gluc = [1, 5, 10, 15, 25, 50]
    Ins_sec = np.zeros((N, len(Gluc)))
    
    
    # calculate mean insulin sectrtion
    for n in range(N):
        if N != 1:
            df = pd.read_csv('/home/phenning/pyABC/Fit_Ins/output/output'+random_string+'_'+ str(n) +'_rule_counts.csv', sep=',')
        else:
            df = pd.read_csv('/home/phenning/pyABC/Fit_Ins/output/output'+random_string+'_rule_counts.csv', sep=',')
        tmp = df['RRP_100:rrp -> Ins_100 @@ if ##(Ca_100 in Cell)<Ca_threshold then kf_ca1*(##(Ca_100 in Cell))*#rrp else kf_ca2*(##(Ca_100 in Cell))*#rrp']
        Ins_sec[n, 0] = (tmp[(np.abs(time-1.75*3600)).argmin()]-tmp[(np.abs(time-0.25*3600)).argmin()])/(time[(np.abs(time-1.75*3600)).argmin()]-time[(np.abs(time-0.25*3600)).argmin()])
        Ins_sec[n, 1] = (tmp[(np.abs(time-3.75*3600)).argmin()]-tmp[(np.abs(time-2.25*3600)).argmin()])/(time[(np.abs(time-3.75*3600)).argmin()]-time[(np.abs(time-2.25*3600)).argmin()])
        Ins_sec[n, 2] = (tmp[(np.abs(time-5.75*3600)).argmin()]-tmp[(np.abs(time-4.25*3600)).argmin()])/(time[(np.abs(time-5.75*3600)).argmin()]-time[(np.abs(time-4.25*3600)).argmin()])
        Ins_sec[n, 3] = (tmp[(np.abs(time-7.75*3600)).argmin()]-tmp[(np.abs(time-6.25*3600)).argmin()])/(time[(np.abs(time-7.75*3600)).argmin()]-time[(np.abs(time-6.25*3600)).argmin()])
        Ins_sec[n, 4] = (tmp[(np.abs(time-9.75*3600)).argmin()]-tmp[(np.abs(time-8.25*3600)).argmin()])/(time[(np.abs(time-9.75*3600)).argmin()]-time[(np.abs(time-8.25*3600)).argmin()])
        Ins_sec[n, 5] = (tmp[(np.abs(time-11.75*3600)).argmin()]-tmp[(np.abs(time-10.25*3600)).argmin()])/(time[(np.abs(time-11.75*3600)).argmin()]-time[(np.abs(time-10.25*3600)).argmin()])

    output = np.mean(Ins_sec, 0)
    
    # deletes output files
    for filename in glob.glob('/home/phenning/pyABC/Fit_Ins/output/*'+random_string+'*'):
        os.remove(filename) 
    for filename in glob.glob('/home/phenning/pyABC/Fit_Ins/*'+random_string+'*'):
        os.remove(filename)
    return{"Ins_sec":output}

# calcules (relative) mean square error
def distance(sim, data):
    error = (
        np.sum(pow((sim['Ins_sec'] - data['Ins_sec'])/data['Ins_sec'],2))
    )
    return error

def run_ABC():
    prior = pyabc.Distribution(
        Ca_threshold=pyabc.RV("uniform", 700, 400),
        kr_gluc=pyabc.RV("uniform", 210, 100),
        kr_offset=pyabc.RV("uniform", 4.3, 1),
        kr_ca=pyabc.RV("uniform", 0.08, 0.04),
        kf_ca1=pyabc.RV("uniform", 0.0005, 0.0015),
        kf_ca2=pyabc.RV("uniform", 0.0015, 0.0035)
    )
    
    abc = pyabc.ABCSMC(
        models=model,
        parameter_priors=prior,
        distance_function=distance,
        population_size=pyabc.AdaptivePopulationSize(1000, 0.14, 2500),
        sampler = pyabc.sampler.MulticoreEvalParallelSampler(n_procs=250),
    )
    
    # Insulin secretion data from Ishihara et al. 1993
    theo = {'Ins_sec': np.array([4.775565227, 6.63200055, 22.21985869, 35.69815105, 46.07523568, 48.20167101])}
    
    import tempfile
    db_path = "sqlite:///" + os.path.join(tempfile.gettempdir(), "/home/phenning/pyABC/Fit_Ins/pyABC_Ins.db")
    abc_id = abc.new(db_path, {"Ins_sec": theo['Ins_sec']})
    
    history = abc.run(minimum_epsilon=0.01, max_nr_populations=4)
    return(abc_id, history, db_path)

def continue_ABC(model, prior, distance, db_path, abc_id):
    abc_continued = pyabc.ABCSMC(
        model,
        prior,
        distance,
        population_size=pyabc.AdaptivePopulationSize(1000, 0.14, 2500),
        sampler = pyabc.sampler.MulticoreEvalParallelSampler(n_procs=250))
    abc_continued.load(db_path, abc_id)
    abc_continued.run(minimum_epsilon=0.01, max_nr_populations=10)
    
    return(abc_continued)
    
def plot_ABC(path, model, distance, id=1):
    prior = pyabc.Distribution(
        Ca_threshold=pyabc.RV("uniform", 700, 400),
        kr_gluc=pyabc.RV("uniform", 210, 100),
        kr_offset=pyabc.RV("uniform", 4.3, 1),
        kr_ca=pyabc.RV("uniform", 0.08, 0.04),
        kf_ca1=pyabc.RV("uniform", 0.0005, 0.0015),
        kf_ca2=pyabc.RV("uniform", 0.0015, 0.0035)
    )
    abc_continued = pyabc.ABCSMC(model, prior, distance)
    abc_continued.load(path, id)
    history = abc_continued.history
    print('max_t = ', history.max_t)
    import matplotlib.pyplot as plt
    from pyabc.visualization import plot_kde_2d
    from pyabc.visualization import plot_kde_matrix
    fig = plt.figure(figsize=(10,8))
    dfw = history.get_distribution(m=0)
    plot_kde_matrix(
        *dfw,
        limits={
		"Ca_threshold":(700, 1100),
		"kr_gluc":(210, 310),
		"kr_offset":(4.3, 5.3),
		"kr_ca":(0.08, 0.12),
		"kf_ca1":(0.0005, 0.002),
		"kf_ca2":(0.0015, 0.005)})
    plt. title("Results from id: "+str(id))
    plt.savefig('k_Ins_id_'+str(id)+'.png')
    
def output_results(db_path):
    history = pyabc.History(db_path)
    df,w = history.get_distribution()
    names = list(df.columns.values)
    for i in names:
        print(i, " = ",pyabc.weighted_statistics.weighted_mean(np.array(df[i]),w))


if __name__ == '__main__':
	# ----- run pyABC ----- 
    abc_id, history, db_path = run_ABC()
    
    # ----- continu pyABC -----   
    #import tempfile
    #db_path = "sqlite:///" + os.path.join(tempfile.gettempdir(), "/home/phenning/pyABC/Fit_Ins/pyABC_Ins.db")
    #prior = pyabc.Distribution(
    #    Ca_threshold=pyabc.RV("uniform", 700, 400),
    #    kr_gluc=pyabc.RV("uniform", 210, 100),
    #    kr_offset=pyabc.RV("uniform", 4.3, 1),
    #    kr_ca=pyabc.RV("uniform", 0.08, 0.04),
    #    kf_ca1=pyabc.RV("uniform", 0.0005, 0.0015),
    #    kf_ca2=pyabc.RV("uniform", 0.0015, 0.0035)
    #)
    #abc_continued = continue_ABC(model = model, prior = prior, distance = distance, db_path = db_path, abc_id = 1)
    
	# ----- plot results -----
    #import tempfile
    #path = "sqlite:///" + os.path.join(tempfile.gettempdir(), "/home/phenning/pyABC/Fit_Ins/pyABC_Ins.db")
    #plot_ABC(path = path, model = model, distance = distance, id = 2)
    
	# ----- output results -----
    #import tempfile
    #db_path = "sqlite:///" + os.path.join(tempfile.gettempdir(), "/home/phenning/pyABC/Fit_Ins/pyABC_Ins.db")
    #output_results(db_path)
    
```
