## Appending and model files. for "Illuminating the Role of Asymmetric Mitochondrial Fission on Beta-Cell Health": S9_file.py

```
# -*- coding: utf-8 -*-
"""
Created on Thu May 16 09:52:16 2024

@author: phenning
"""

import numpy as np
import os
import pandas as pd
from timeit import default_timer as timer
import subprocess
import pyabc
import random
import string
import glob
import matplotlib.pyplot as plt


# def simulation model
baseline = """
/* cAMP submodel */

/* ----- rate constants ----- */
const vol_cell = 1025 [micro^3 m^3];
const NA = 6.022e23 [1/mol];

const Vm_glyc = 3408.964 [1/min];
const Km_glyc = 4.626 [milli mol/L];
const Ki_glyc = 42806.181;

const Vmax_LDH = 5450 [1/min];
const Km_LDH_pyr = 47.5 [micro mol/L]*vol_cell*NA;

const Vmax_PDH = 363.333 [1/min];
const Km_PDH_pyr = 47.5 [micro mol/L]*vol_cell*NA;

const vol_total = 75 [micro^3 m^3];

const k_ATP_u = 0.748 [1/min];

/* Ca2+ dynamic */
const Ca_min = 486.4;
const Ca_diff = 742.5;
const Ca_rise = 0.000759;
const Ca_ATP = 8719;
const k_ca_flux = 10 [1/s];

/* cAMP dynamic */
const k_cAMP_d = [1/min];
const vmax_AC = 8000 [1/min];
const Km_Ca = ;

/* ----- Species ----- */
Cell(Gluc: float[mol/L]);
Mito();
pyruvat_1000000();
ATP_1000000();
Ca_100();
cAMP_100();

/* ----- Initial solution ----- */
Initial: 1 Cell(Gluc = 2[milli mol/L]){75 Mito + 4000 ATP_1000000 + 400 Ca_100 + 6200 cAMP_100};

/* ----- Rules ----- */
/* glycolysis of 1000000 glucose molecule*/
Cell{ATP_1000000:atp} -> Cell{2 pyruvat_1000000 + 3 ATP_1000000} @@ Vm_glyc*Cell.Gluc/((Km_glyc+Cell.Gluc)*(1+#atp/Ki_glyc));

/* anerobmic fermentaion */
pyruvat_1000000:pyr -> @@ Vmax_LDH*#pyr*1e6/(#pyr*1e6+Km_LDH_pyr);

/* TCA + OxPhos*/
Mito + pyruvat_1000000:pyr -> Mito + 15 ATP_1000000 @@ (1/75)*Vmax_PDH*#pyr*1e6/(#pyr*1e6+Km_PDH_pyr);

/* ATP usage */
ATP_1000000:atp ->  @@ k_ATP_u*#atp;

/* Ca2+ influx */
Cell{Ca_100:ca+ATP_1000000:atp} -> Cell{Ca_100+Ca_100+ATP_1000000} @@ if (Ca_min + Ca_diff/(1+exp(Ca_rise*(-1*(#atp)+Ca_ATP))))>#ca then k_ca_flux;

/* Ca2+ outflux */
Cell{Ca_100:ca+ATP_1000000:atp} -> Cell{ATP_1000000} @@ if (Ca_min + Ca_diff/(1+exp(Ca_rise*(-1*(#atp)+Ca_ATP))))<#ca then k_ca_flux;

/* -- cAMP Dynamic -- */
/* cAMP reation (dynamics of sAC: Litvin et al. 2003, dynamic of AC1: Gille 2004 et al.) */
/* calcium regulation taken from Peercy et al. 2015 */
Cell -> Cell{cAMP_100} @@ vmax_AC*((##(Ca_100 in Cell))^2)/(Km_Ca^2+((##(Ca_100 in Cell))^2));

/* decay of cAMP */
Cell{cAMP_100:c} -> Cell @ k_cAMP_d;

/* ----- Glucose rise ------ */

Cell -> Cell(Gluc = 20 [milli mol/L]) @@ if time() > 2[h] && time() < 2.1[h] && Cell.Gluc<20 [milli mol/L] then immediately;

output ##(cAMP_100);
"""

# added rates to the base model and execute it
def model(rates, t_end=4, Nt=150, N=1):
    silent = True
    
    # adds fit parameter to basemodel    
    k_cAMP_d = rates['k_cAMP_d']
    if k_cAMP_d == None:
        raise ValueError('No k_cAMP_d specified')
    model = baseline.replace("const k_cAMP_d = ","const k_cAMP_d = " + str(k_cAMP_d))
        
    Km_Ca = rates['Km_Ca']
    if Km_Ca == None:
        raise ValueError('Km_Ca specified')
    model = model.replace("const Km_Ca = ","const Km_Ca = " + str(Km_Ca))
      
    # creates model file with random name
    random_string = "_"+''.join(random.choice(string.ascii_lowercase) for i in range(25))
    text_file = open("/home/phenning/pyABC/Fit_cAMP/run_3/model"+random_string+".mlr", "w")
    text_file.write(model)
    text_file.close()
    
    # runs ML-Rules3 model
    if silent:
            subprocess.check_output('cargo run --release --manifest-path /home/phenning/ml-rules-3/Cargo.toml -- /home/phenning/pyABC/Fit_cAMP/run_3/model'+random_string+'.mlr --simulate-until "'+str(t_end)+'[h]" --num-observation '+str(Nt)+' --dynamic-silent --output-prefix /home/phenning/pyABC/Fit_cAMP/run_3/output/output'+random_string+' --num-reps '+str(N)+' --output-csv', shell=True,stderr=subprocess.STDOUT)
    else:
        subprocess.check_output('cargo run --release --manifest-path /home/phenning/ml-rules-3/Cargo.toml -- /home/phenning/pyABC/Fit_cAMP/run_3/model'+random_string+'.mlr --simulate-until "'+str(t_end)+'[h]" --num-observation '+str(Nt)+' --dynamic-silent --output-prefix /home/phenning/pyABC/Fit_cAMP/run_3/output/output'+random_string+' --num-reps '+str(N)+' --output-csv', shell=True)

    # load output data
    if N != 1:
        df = pd.read_csv('/home/phenning/pyABC/Fit_cAMP/run_3/output/output'+random_string+'_0.csv', sep=',')
    else:
        df = pd.read_csv('/home/phenning/pyABC/Fit_cAMP/run_3/output/output'+random_string+'.csv', sep=',')
    time = np.unique(df['time[s]'])
    
    Gluc = [2, 20]
    cAMP = np.zeros((len(Gluc), N))
    
    
    # calculate mean cAMP level
    for n in range(N):
        if N != 1:
            df = pd.read_csv('/home/phenning/pyABC/Fit_cAMP/run_3/output/output'+random_string+'_'+ str(n) +'.csv', sep=',')
        else:
            df = pd.read_csv('/home/phenning/pyABC/Fit_cAMP/run_3/output/output'+random_string+'.csv', sep=',')
        tmp = df['##(cAMP_100)']
        for i in range(len(Gluc)):
            cAMP[i,n] = (np.mean(tmp[np.logical_and(time>3600*(2*i+0.25), time<3600*(2*i+1.75))]))
        output = np.mean(cAMP, 1)
    
    # deletes output files
    for filename in glob.glob('/home/phenning/pyABC/Fit_cAMP/run_3/output/*'+random_string+'*'):
        os.remove(filename) 
    for filename in glob.glob('/home/phenning/pyABC/Fit_cAMP/run_3/*'+random_string+'*'):
        os.remove(filename)
    return{"cAMP":output}

# calcules (relative) mean square error
def distance(sim, data):
    error = (
        np.sum(pow((sim['cAMP'] - data['cAMP'])/data['cAMP'],2))
    )
    return error

def run_ABC():
    prior = pyabc.Distribution(
        k_cAMP_d = pyabc.RV("uniform", 0.2, 0.8),
        Km_Ca = pyabc.RV("uniform", 500, 1500),
        )
    
    abc = pyabc.ABCSMC(
        models=model,
        parameter_priors=prior,
        distance_function=distance,
        population_size=pyabc.AdaptivePopulationSize(750, 0.14, 2500),
        sampler = pyabc.sampler.MulticoreEvalParallelSampler(n_procs=126),
    )
    
    # cAMP data from Jing et al. 2021
    theo = {'cAMP': np.array([621896.8435, 1681807.411])/100}
    
    import tempfile
    db_path = "sqlite:///" + os.path.join(tempfile.gettempdir(), "/home/phenning/pyABC/Fit_cAMP/run_3/pyABC_cAMP.db")
    abc_id = abc.new(db_path, {"cAMP": theo['cAMP']})
    
    history = abc.run(minimum_epsilon=0.01, max_nr_populations=10)
    return(abc_id, history, db_path)

def continue_ABC(model, prior, distance, db_path, abc_id):
    abc_continued = pyabc.ABCSMC(
        model,
        prior,
        distance,
        population_size=pyabc.AdaptivePopulationSize(750, 0.14, 2500),
        sampler = pyabc.sampler.MulticoreEvalParallelSampler(n_procs=126))
    abc_continued.load(db_path, abc_id)
    abc_continued.run(minimum_epsilon=0.001, max_nr_populations=10)
    
    return(abc_continued)
    
def plot_ABC(path, model, distance, id=1):
    prior = pyabc.Distribution(
        k_cAMP_d = pyabc.RV("uniform", 0.2, 0.8),
        Km_Ca = pyabc.RV("uniform", 500, 1500),
        )
    abc_continued = pyabc.ABCSMC(model, prior, distance)
    abc_continued.load(path, id)
    history = abc_continued.history
    print('max_t = ', history.max_t)
    import matplotlib.pyplot as plt
    from pyabc.visualization import plot_kde_2d
    from pyabc.visualization import plot_kde_matrix
    fig, ax = plt.subplots()
    dfw = history.get_distribution(m=0)
    plot_kde_matrix(
        *dfw,
        limits={
        	"k_cAMP_d":(0.2, 1),
                "Km_Ca": (500, 2000)}
        )
    plt. title("Results from id: "+str(id))
    plt.savefig('k_cAMP_id_'+str(id)+'.png')
    
def output_results(db_path):
    history = pyabc.History(db_path)
    df,w = history.get_distribution()
    names = list(df.columns.values)
    print('Weighted mean:')
    for i in names:
        print(i, " = ",pyabc.weighted_statistics.weighted_mean(np.array(df[i]),w))
    print('Weighted median:')
    for i in names:
        print(i, " = ",pyabc.weighted_statistics.weighted_median(np.array(df[i]),w))


if __name__ == '__main__':
	# ----- run pyABC -----
    abc_id, history, db_path = run_ABC()
    
    # ----- continu pyABC -----   
    #import tempfile
    #db_path = "sqlite:///" + os.path.join(tempfile.gettempdir(), "/home/phenning/pyABC/Fit_cAMP/run_3/pyABC_cAMP.db")
    #prior = pyabc.Distribution(
    #    k_cAMP_d = pyabc.RV("uniform", 0.001, 0.5),
    #    n = pyabc.RV("uniform", 1, 4),
    #    )
    #abc_continued = continue_ABC(model = model, prior = prior, distance = distance, db_path = db_path, abc_id = 1)
    
	# ----- plot results -----
    #import tempfile
    #path = "sqlite:///" + os.path.join(tempfile.gettempdir(), "/home/phenning/pyABC/Fit_cAMP/run_3/pyABC_cAMP.db")
    #plot_ABC(path = path, model = model, distance = distance, id = 21)
    
	# ----- output results -----
    #import tempfile
    #db_path = "sqlite:///" + os.path.join(tempfile.gettempdir(), "/home/phenning/pyABC/Fit_cAMP/run_3/pyABC_cAMP.db")
    #output_results(db_path)
    
```
