## Appending and model files. for "Illuminating the Role of Asymmetric Mitochondrial Fission on Beta-Cell Health": S10_file.py

```
# -*- coding: utf-8 -*-
"""
Created on Thu May 16 09:52:16 2024

@author: phenning
"""

import numpy as np
import os
import pandas as pd
from timeit import default_timer as timer
import subprocess
import random
import string
import glob
import matplotlib.pyplot as plt
#import pyswarms as ps
from pyswarms.single.global_best import GlobalBestPSO
from multiprocessing import Pool
from pyswarms.utils.plotters import (plot_cost_history, plot_contour, plot_surface)

# def simulation model
baseline = """
/* beta-cell model */

/* ----- rate constants ----- */
/* general constants */
const vol_cell = 1025 [micro^3 m^3];
const NA = 6.022e23 [1/mol];

/* fission-fusion dynamic */
const kon = 0.025 [micro^3 m^3/s];
const Kd = 0.301 [micro mol/L]*NA;
const koff = kon*Kd;
const k_fission = 8.211e-08 [Hz];
const N_fission = 11;
const asym_fission = XX;
const k_fusion = 5.41e-4[1/min];

/* CaN dynamic */
const kcatCaN = 7.6 [1/s];
const KMCaN = 160 [micro mol/L];
const K_act_CaN = 14e-9[mol/L]*vol_cell*NA;

/* PKA dynamic */
const kcatpka= 4 [1/s];
const KMpka = 70 [micro mol/L];
const K_act_PKA = 0.9e-6[mol/L]*vol_cell*NA;

/* respiration */
const Vm_glyc = 3408.964 [1/min];
const Km_glyc = 4.626 [milli mol/L];
const Ki_glyc = 42806.181;
const Vmax_LDH = 5450 [1/min];
const Km_LDH_pyr = 47.5 [micro mol/L]*vol_cell*NA;
const Vmax_PDH = 363.333 [1/min];
const Km_PDH_pyr = 47.5 [micro mol/L]*vol_cell*NA;
const f_ATP = XX;
const damage_lim = XX;
const damage_m = XX;

const k_ATP_u = 0.748 [1/min];

/* Ca2+ dynamic */
const Ca_min = 486.4;
const Ca_diff = 742.5;
const Ca_rise = 0.000759;
const Ca_ATP = 8719;
const k_ca_flux = 10 [1/s];

/* Insulin dynamic */
const Ca_threshold = 852.936;
const kr_gluc = 261.542 [L/mol s];
const kr_offset = 4.796 [Hz];
const kr_ca = 0.103 [1/s];
const kf_ca1 = 0.00125 [1/s];
const kf_ca2 = 0.00326 [1/s];

/* cAMP dynamic */
const vmax_AC = 8000 [1/min];
const k_cAMP_d = 0.2602 [1/min];
const Km_Ca = 1045;

/* biogenesis */
const N0 = 6800;
const gamma = 15;
const k_biog = 0.05 [1/day];
const k_repair = 0.1 [1/day];

const Thr_damage = XX;


/* ----- Species ----- */
Cell(Gluc: float[mol/L], mito_vol:int*);
Mito(vol:int*, health:int*, damage:int*, rand1:float*, rand2:float*);
Drp1_10(p: bool);
Fis1();
MFF();
PKA();
cAMP_100();
CaN();
Ca_100();
pyruvat_1000000();
ATP_1000000();
RRP_100();
Ins_100();


/* ----- Initial solution ----- */
Initial: 1 Cell(Gluc = 1 [milli mol/L], mito_vol=6800){68 Mito(vol = 100, health=XX, damage=XX, rand1=1001, rand2=1001) + 10000 Drp1_10(p=false) + 10000 Drp1_10(p=true) + 300000 PKA() + 350000 CaN() + 4000 ATP_1000000 + 500 Ca_100 + 6200 cAMP_100 + 306 Fis1() + 306 MFF()};

/* ----- Rules ----- */
/* Drp1_10 binding */
Mito{Fis1} + Drp1_10(p==false) -> Mito{Fis1{Drp1_10}} @ kon/vol_cell;
Mito{MFF} + Drp1_10(p==false) -> Mito{MFF{Drp1_10}} @ kon/vol_cell;

/* Drp1_10 unbinding */
Mito{Fis1:f{Drp1_10}} -> Mito{Fis1} + Drp1_10 @@ 1/10*koff*#f;
Mito{MFF:m{Drp1_10}} -> Mito{MFF} + Drp1_10 @@ 1/10*koff*#m;

/* phosphorylation by PKA */
Cell{Drp1_10(p==false):d + PKA:p} -> Cell{Drp1_10(p=true) + PKA} @@ 1/10*kcatpka*#p*#d*10/(KMpka*vol_cell*NA+#d*10)*100*##(cAMP_100 in Cell)/(K_act_PKA+100*##(cAMP_100 in Cell));

/* dephosphorylation by CaN */
Cell{Drp1_10(p==true):d + CaN:c} -> Cell{Drp1_10(p=false) + CaN} @@ 1/10*kcatCaN*#c*#d*10/(KMCaN*vol_cell*NA+#d*10)*100*##(Ca_100 in Cell)/(K_act_CaN+100*##(Ca_100 in Cell));


/* Midpoint fission */
// generate random number for fission event
Mito:m{MFF{Drp1_10:drp + ?solA}} -> Mito(rand1 = RandomGauss(0.5, 0.075)){MFF}+Drp1_10+?solA @@ if (#drp>N_fission) then k_fission*(m.vol)^3;

// executed fission event
Mito(rand1<1000):m{?sol1[Mito.rand1] + ?sol2[1-Mito.rand1]} -> Mito(vol=ceil(m.vol*m.rand1), health=ceil(m.health*m.rand1), damage=ceil(m.damage*m.rand1), rand1=1001, rand2=1001){?sol1} + Mito(vol=floor(m.vol*(1-m.rand1)), health = floor(m.health*(1-m.rand1)), damage = floor(m.damage*(1-m.rand1)), rand1=1001, rand2=1001){?sol2} @@ immediately;


/* Peripheral Fission*/
// generate random number for fission event
Mito:m{Fis1{Drp1_10:drp + ?solA}} -> Mito(rand2 = RandomGauss(0.2, 0.04)){Fis1}+Drp1_10+?solA @@ if (#drp>N_fission) then k_fission*(m.vol)^3;

// executed fission event
/* Damage Selective Fission */
Mito(rand2<1000, rand2>asym_fission, health+damage>=2):m{?sol1[Mito.rand2] + ?sol2[1-Mito.rand2]} -> Mito(vol=ceil(m.vol*m.rand2), health=ceil(m.health*m.rand2-m.damage*asym_fission), damage=ceil(m.damage*(m.rand2+asym_fission)), rand1=1001, rand2=1001){?sol1} + Mito(vol=floor(m.vol*(1-m.rand2)), health=floor(m.health*(1-m.rand2)+m.damage*asym_fission), damage=floor(m.damage*((1-m.rand2)-asym_fission)), rand1=1001, rand2=1001){?sol2} @@ immediately;

// checks that random number would not create negative volume or health values
Mito(rand2<=asym_fission) -> Mito(rand2=asym_fission + 0.0001) @@ immediately;


/* fusion */
Mito:m1{?solm1} + Mito:m2{?solm2} -> Mito(vol=m1.vol+m2.vol, health=m1.health+m2.health, damage=m1.damage+m2.damage, rand1=1001, rand2=1001){?solm1 + ?solm2} @@ if ((m1.damage/(m1.health+m1.damage))<Thr_damage && (m2.damage/(m2.health+m2.damage))<Thr_damage) then k_fusion;

/* glycolysis of 1000000 glucose molecule*/
Cell{ATP_1000000:atp} -> Cell{2 pyruvat_1000000 + 3 ATP_1000000} @@ Vm_glyc*Cell.Gluc/((Km_glyc+Cell.Gluc)*(1+#atp/Ki_glyc));

/* anerobmic fermentaion */
pyruvat_1000000:pyr -> @@ Vmax_LDH*#pyr*1e6/(#pyr*1e6+Km_LDH_pyr);

/* TCA + OxPhos*/
Mito:m + pyruvat_1000000:pyr -> Mito + 15 ATP_1000000 @@
f_ATP*(0.1+0.9/(1+exp(damage_m*(damage_lim-m.health/(m.health+m.damage)))))*(m.vol/N0)*Vmax_PDH*#pyr*1e6/(#pyr*1e6+Km_PDH_pyr);

/* ATP usage */
ATP_1000000:atp ->  @@ k_ATP_u*#atp;

/* Ca2+ influx */
Cell{Ca_100:ca+ATP_1000000:atp} -> Cell{Ca_100+Ca_100+ATP_1000000} @@ if (Ca_min + Ca_diff/(1+exp(Ca_rise*(-1*(#atp)+Ca_ATP))))>#ca then k_ca_flux;

/* Ca2+ outflux */
Cell{Ca_100:ca+ATP_1000000:atp} -> Cell{ATP_1000000} @@ if (Ca_min + Ca_diff/(1+exp(Ca_rise*(-1*(#atp)+Ca_ATP))))<#ca then k_ca_flux;

/* cAMP Dynamic */
/* cAMP reation (dynamics of sAC: Litvin et al. 2003, dynamic of AC1: Gille 2004 et al.) */
/* calcium regulation taken from Peercy et al. 2015 */
Cell -> Cell{cAMP_100} @@ vmax_AC*((##(Ca_100 in Cell))^2)/(Km_Ca^2+((##(Ca_100 in Cell))^2));

/* decay of cAMP */
Cell{cAMP_100:c} -> Cell @ k_cAMP_d;

/* RRP + Insulin model taken from Grespan et al. 2017 */
/* filling of the readily releasable pool (RRL) */
Cell -> Cell{RRP_100} @@ if ##(Ca_100 in Cell)<Ca_threshold then kr_gluc*Cell.Gluc+kr_offset else kr_gluc*Cell.Gluc+kr_offset+kr_ca*(##(Ca_100 in Cell)-Ca_threshold);

/* Insulin secretion from RRP */
RRP_100:rrp -> Ins_100 @@ if ##(Ca_100 in Cell)<Ca_threshold then kf_ca1*(##(Ca_100 in Cell))*#rrp else kf_ca2*(##(Ca_100 in Cell))*#rrp;

/* damage by oxPhos */
Mito(health>0):m + pyruvat_1000000:pyr -> Mito(damage = m.damage+1, health = m.health-1) + pyruvat_1000000 @@ 
f_ATP*(1+0.5*m.damage/(m.health+m.damage))*(0.1+0.9/(1+exp(damage_m*(damage_lim-m.health/(m.health+m.damage)))))*(m.vol/N0)*0.007*Vmax_PDH*#pyr*1e6/(#pyr*1e6+Km_PDH_pyr);

/* damage repair */
Mito(damage>0):m -> Mito(damage = m.damage-1, health = m.health+1) @@ k_repair*m.damage;

/* mitophagy */
Cell:c{Mito:m{?solm}} -> Cell(mito_vol=c.mito_vol-m.vol){?solm} @@ if (m.vol < 50 && m.damage/(m.damage+m.health) >= Thr_damage) then 10 [Hz];

/* Deal with species after removal */
Cell{Mito:m + Fis1} -> Cell{Mito{Fis1}} @@ 1000 [Hz]*(m.vol);
Cell{Mito:m + MFF} -> Cell{Mito{MFF}} @@ 1000 [Hz]*(m.vol);

/* biogenesis */
Cell:c{Mito:m} -> Cell(mito_vol=c.mito_vol+1){Mito(vol = m.vol+1, health = m.health+1)} @@ k_biog*N0/(1+exp((c.mito_vol-(N0-100))/gamma));

/* ----- Glucose rise ------ */

Cell -> Cell(Gluc = 5 [milli mol/L]) @@ if time() > 10[h] && time() < 10.1[h] && Cell.Gluc<5 [milli mol/L] then immediately;
Cell -> Cell(Gluc = 10 [milli mol/L]) @@ if time() > 16[h] && time() < 16.1[h] && Cell.Gluc<10 [milli mol/L] then immediately;
Cell -> Cell(Gluc = 15 [milli mol/L]) @@ if time() > 22[h] && time() < 22.1[h] && Cell.Gluc<15 [milli mol/L] then immediately;
Cell -> Cell(Gluc = 25 [milli mol/L]) @@ if time() > 28[h] && time() < 28.1[h] && Cell.Gluc<25 [milli mol/L] then immediately;
Cell -> Cell(Gluc = 50 [milli mol/L]) @@ if time() > 34[h] && time() < 34.1[h] && Cell.Gluc<50 [milli mol/L] then immediately;
"""

# executes model in parallel
def run_par(rates):
    rates = rates.tolist()
    results = []
    pool = Pool()
    res = pool.map(model, rates)
    results.append(res)
    pool.close()
    pool.join()
    return np.array(results[0])

# exectes model one at a time
def run_seq(rates):
    n_particles = rates.shape[0]
    j = [model(rates[i,:]) for i in range(n_particles)]
    return np.array(j)

# added rates to the base model and execute it
def model(rates, t_end_base=40, t_end_Fis1=12, Nt=300, N=2):
    silent = True
    
    # adds fit parameter to basemodel
    damage_lim = float(rates[0])
    if damage_lim == None:
        raise ValueError('No damage_lim specified')
    model = baseline.replace("const damage_lim = XX","const damage_lim = " + str(damage_lim))
    
    damage_m = float(rates[1])
    if damage_m == None:
        raise ValueError('No damage_m specified')
    model = model.replace("const damage_m = XX","const damage_m = " + str(damage_m))
    
    Thr_damage = float(rates[2])
    if Thr_damage == None:
        raise ValueError('No Thr_damage specified')
    model = model.replace("const Thr_damage = XX","const Thr_damage = " + str(Thr_damage))

    model = model.replace("health=XX","health=" + str(100-round(Thr_damage*100)+5))
    model = model.replace("damage=XX","damage=" + str(round(Thr_damage*100)-5))
    
    asym_fission = float(rates[3])
    if asym_fission == None:
        raise ValueError('No asym_fission specified')
    model = model.replace("const asym_fission = XX","const asym_fission = " + str(asym_fission))
    
    f_ATP = float(rates[4])
    if f_ATP == None:
        raise ValueError('No f_ATP specified')
    model = model.replace("const f_ATP = XX","const f_ATP = " + str(f_ATP))

        
    # creates model file with random name
    random_string = "_"+''.join(random.choice(string.ascii_lowercase) for i in range(25))
    text_file = open("/home/phenning/pySwarm/Fit_damage_response/model_base"+random_string+".mlr", "w")
    text_file.write(model)
    text_file.close()

    Fis = 48
    if Fis == None:
        raise ValueError('No Fis specified')
    model_Fis = model.replace("306 Fis1() +",str(Fis) + " Fis1() +")

    model_Fis = model_Fis.replace("Cell(Gluc = 1 [milli mol/L]", "Cell(Gluc = 25 [milli mol/L]")
    model_Fis = model_Fis.replace("Cell -> Cell(Gluc = 5 [milli mol/L]) @@ if time() > 10[h] && time() < 10.1[h] && Cell.Gluc<5 [milli mol/L] then immediately;", " ")
    model_Fis = model_Fis.replace("Cell -> Cell(Gluc = 10 [milli mol/L]) @@ if time() > 16[h] && time() < 16.1[h] && Cell.Gluc<10 [milli mol/L] then immediately;", " ")
    model_Fis = model_Fis.replace("Cell -> Cell(Gluc = 15 [milli mol/L]) @@ if time() > 22[h] && time() < 22.1[h] && Cell.Gluc<15 [milli mol/L] then immediately;", " ")
    model_Fis = model_Fis.replace("Cell -> Cell(Gluc = 25 [milli mol/L]) @@ if time() > 28[h] && time() < 28.1[h] && Cell.Gluc<25 [milli mol/L] then immediately;", " ")
    model_Fis = model_Fis.replace("Cell -> Cell(Gluc = 50 [milli mol/L]) @@ if time() > 34[h] && time() < 34.1[h] && Cell.Gluc<50 [milli mol/L] then immediately;", " ")

    text_file = open("/home/phenning/pySwarm/Fit_damage_response/model_Fis"+random_string+".mlr", "w")
    text_file.write(model_Fis)
    text_file.close()

    # runs ML-Rules3 model

    if silent:
        subprocess.check_output('cargo run --release --manifest-path /home/phenning/ml-rules-3/Cargo.toml -- /home/phenning/pySwarm/Fit_damage_response/model_base'+random_string+'.mlr --simulate-until "'+str(t_end_base)+'[h]" --num-observation '+str(t_end_base*10)+' --dynamic-silent --output-prefix /home/phenning/pySwarm/Fit_damage_response/output/output_base'+random_string+' --num-reps '+str(N)+' --output-flat-table --output-rule-counts --output-csv', shell=True,stderr=subprocess.STDOUT)
    else:
        subprocess.check_output('cargo run --release --manifest-path /home/phenning/ml-rules-3/Cargo.toml -- /home/phenning/pySwarm/Fit_damage_response/model_base'+random_string+'.mlr --simulate-until "'+str(t_end_base)+'[h]" --num-observation '+str(t_end_base*10)+' --dynamic-silent --output-prefix /home/phenning/pySwarm/Fit_damage_response/output/output_base'+random_string+' --num-reps '+str(N)+' --output-flat-table --output-rule-counts --output-csv', shell=True)

    if silent:
        subprocess.check_output('cargo run --release --manifest-path /home/phenning/ml-rules-3/Cargo.toml -- /home/phenning/pySwarm/Fit_damage_response/model_Fis'+random_string+'.mlr --simulate-until "'+str(t_end_Fis1)+'[h]" --num-observation '+str(t_end_Fis1*10)+' --dynamic-silent --output-prefix /home/phenning/pySwarm/Fit_damage_response/output/output_Fis'+random_string+' --num-reps '+str(N)+' --output-flat-table --output-rule-counts --output-csv', shell=True,stderr=subprocess.STDOUT)
    else:
        subprocess.check_output('cargo run --release --manifest-path /home/phenning/ml-rules-3/Cargo.toml -- /home/phenning/pySwarm/Fit_damage_response/model_Fis'+random_string+'.mlr --simulate-until "'+str(t_end_Fis1)+'[h]" --num-observation '+str(t_end_Fis1*10)+' --dynamic-silent --output-prefix /home/phenning/pySwarm/Fit_damage_response/output/output_Fis'+random_string+' --num-reps '+str(N)+' --output-flat-table --output-rule-counts --output-csv', shell=True)

    # load output data
    if N != 1:
        df = pd.read_csv('/home/phenning/pySwarm/Fit_damage_response/output/output_base'+random_string+'_0.csv', sep=',')
    else:
        df = pd.read_csv('/home/phenning/pySwarm/Fit_damage_response/output/output_base'+random_string+'.csv', sep=',')
    time = np.unique(df['time[s]'])
    
    Gluc = [1, 5, 10, 15, 25, 50]
    t = [10, 14, 18, 22, 26, 30]
    Ins_sec_base = np.zeros((N, len(Gluc)))
    Ins_sec_Fis = np.zeros((N, len(Gluc)))

    # calculate mean insulin sectrtion
    for n in range(N):
        if N != 1:
            df = pd.read_csv('/home/phenning/pySwarm/Fit_damage_response/output/output_base'+random_string+'_'+ str(n) +'_rule_counts.csv', sep=',')
        else:
            df = pd.read_csv('/home/phenning/pySwarm/Fit_damage_response/output/output_base'+random_string+'_rule_counts.csv', sep=',')
        tmp = df['RRP_100:rrp -> Ins_100 @@ if ##(Ca_100 in Cell)<Ca_threshold then kf_ca1*(##(Ca_100 in Cell))*#rrp else kf_ca2*(##(Ca_100 in Cell))*#rrp']
        Ins_sec_base[n, 0] = (tmp[(np.abs(time-t[0]*3600)).argmin()]-tmp[(np.abs(time-(t[0]-2)*3600)).argmin()])/(time[(np.abs(time-t[0]*3600)).argmin()]-time[(np.abs(time-(t[0]-2)*3600)).argmin()])
        Ins_sec_base[n, 1] = (tmp[(np.abs(time-t[1]*3600)).argmin()]-tmp[(np.abs(time-(t[1]-2)*3600)).argmin()])/(time[(np.abs(time-t[1]*3600)).argmin()]-time[(np.abs(time-(t[1]-2)*3600)).argmin()])
        Ins_sec_base[n, 2] = (tmp[(np.abs(time-t[2]*3600)).argmin()]-tmp[(np.abs(time-(t[2]-2)*3600)).argmin()])/(time[(np.abs(time-t[2]*3600)).argmin()]-time[(np.abs(time-(t[2]-2)*3600)).argmin()])
        Ins_sec_base[n, 3] = (tmp[(np.abs(time-t[3]*3600)).argmin()]-tmp[(np.abs(time-(t[3]-2)*3600)).argmin()])/(time[(np.abs(time-t[3]*3600)).argmin()]-time[(np.abs(time-(t[3]-2)*3600)).argmin()])
        Ins_sec_base[n, 4] = (tmp[(np.abs(time-t[4]*3600)).argmin()]-tmp[(np.abs(time-(t[4]-2)*3600)).argmin()])/(time[(np.abs(time-t[4]*3600)).argmin()]-time[(np.abs(time-(t[4]-2)*3600)).argmin()])
        Ins_sec_base[n, 5] = (tmp[(np.abs(time-t[5]*3600)).argmin()]-tmp[(np.abs(time-(t[5]-2)*3600)).argmin()])/(time[(np.abs(time-t[5]*3600)).argmin()]-time[(np.abs(time-(t[5]-2)*3600)).argmin()])


    for n in range(N):
        if N != 1:
            df = pd.read_csv('/home/phenning/pySwarm/Fit_damage_response/output/output_Fis'+random_string+'_'+ str(n) +'_rule_counts.csv', sep=',')
        else:
            df = pd.read_csv('/home/phenning/pySwarm/Fit_damage_response/output/output_Fis'+random_string+'_rule_counts.csv', sep=',')
        tmp = df['RRP_100:rrp -> Ins_100 @@ if ##(Ca_100 in Cell)<Ca_threshold then kf_ca1*(##(Ca_100 in Cell))*#rrp else kf_ca2*(##(Ca_100 in Cell))*#rrp']
        Ins_sec_Fis[n, 0] = (tmp[(np.abs(time-t_end_Fis1*3600)).argmin()]-tmp[(np.abs(time-(t_end_Fis1-2)*3600)).argmin()])/(time[(np.abs(time-t_end_Fis1*3600)).argmin()]-time[(np.abs(time-(t_end_Fis1-2)*3600)).argmin()])

    output_base = np.mean(Ins_sec_base, 0)
    output_Fis = np.mean(Ins_sec_Fis, 0)
    
	# Insulin secretion data from Ishihara et al. 1993
	# Insulin secretion at Fis1 KD is reduced by 40% at 25 mM glucose (Twig et al. 2008)
    data = {'Ins_sec_base': np.array([4.775565227, 6.63200055, 22.21985869, 35.69815105, 46.07523568, 48.20167101]), 'Ins_sec_Fis': np.array([46.07523568*0.60])}
    
	# calcules (relative) mean square error
	error = (
        np.sum(pow((output_base - data['Ins_sec_base'])/data['Ins_sec_base'],2) + 6*pow((output_Fis - data['Ins_sec_Fis'])/data['Ins_sec_Fis'],2))
    )

    # deletes output files
    f = open('/home/phenning/pySwarm/Fit_damage_response/best_tmp.txt', "r")
    best_tmp = f.read().split(", ")
    best = float(best_tmp[0])
    best_name = best_tmp[1]
    f.close()
    if error < best:
        for filename in glob.glob('/home/phenning/pySwarm/Fit_damage_response/output/*'+best_name+'*'):
            os.remove(filename) 
        for filename in glob.glob('/home/phenning/pySwarm/Fit_damage_response/*'+best_name+'*'):
            os.remove(filename)
        f = open('/home/phenning/pySwarm/Fit_damage_response/best_tmp.txt', "w")
        f.write(str(error)+', '+random_string)
        f.close()
    else:
        for filename in glob.glob('/home/phenning/pySwarm/Fit_damage_response/output/*'+random_string+'*'):
            os.remove(filename) 
        for filename in glob.glob('/home/phenning/pySwarm/Fit_damage_response/*'+random_string+'*'):
            os.remove(filename)
    return error
    
    
if __name__ == '__main__':
	# Saves the current best configuration for figure plotting
    best = 10000
    best_name = 'abcdefg'
    text_file = open("/home/phenning/pySwarm/Fit_damage_response/best_tmp.txt", "w")
    text_file.write(str(best)+', '+best_name)
    text_file.close()

    # [damage_lim, damage_m, Thr_damage, asym_fission, f_ATP]
    min_bound = np.array([0.25, 10, 0.1, 0, 0.9])
    max_bound = np.array([0.75, 50, 0.9, 0.1, 1.6])
    bounds = (min_bound, max_bound)

    options = {'c1': 0.5, 'c2': 0.3, 'w': 0.9}
    optimizer = GlobalBestPSO(n_particles=125, dimensions=5, options=options, bounds=bounds, oh_strategy={"w":'exp_decay', 'c1':'lin_variation'})
    cost, pos = optimizer.optimize(run_par, 15)
    
    parameter = ['damage_lim', 'damage_m', 'Thr_damage', 'asym_fission', 'f_ATP']
    num = '1'
    log = open("/home/phenning/pySwarm/Fit_damage_response/log_"+str(num)+".txt", "w")
    for i in range(0, len(pos)):
        log.write(str(parameter[i]) + " = " + str(pos[i]) + ", min = " + str(min_bound[i]) + ", max = " + str(max_bound[i]) + "\n")
    log.write("final cost value: " + str(cost))
    log.close()
    
    plot_cost_history(cost_history=optimizer.cost_history)
    plt.savefig('cost_damage_fit_'+str(num)+'.png')
    print('best fit in:'+best_name)
```
